# Structural basis of K^+^/H^+^ antiport in YcgO and its inhibition by unphosphorylated PtsN

**DOI:** 10.64898/2026.08.14.744764

**Authors:** Ayushi Srivastava, Arunabh Athreya, Yogesh Patidar, Vijeta Singh, Abhijit A. Sardesai, Aravind Penmatsa

## Abstract

Cation-proton antiporters (CPAs) are vital for the maintenance of ionic homeostasis and normal physiology among diverse cell types. Despite recent insights into K^+^/H^+^ exchange transporters, the diversity in their structural organization and regulatory mechanisms of K^+^-specific CPAs are minimally understood. Here, we explore the architecture of an *E. coli* CPA1 K^+^/H^+^ antiporter, YcgO and its inhibition by the unphosphorylated form of PtsN, the terminal protein of a regulatory phosphorelay, using cryoEM structures at 3.4 Å and 3.2 Å resolution, respectively. Homodimeric YcgO bound to K^+^ ions in the occluded conformation, harbors additional linked cytosolic domains, RCK and CorC, to regulate the movement of the transport helices within the YcgO dimer. These domains are the sites of interaction and efflux inhibition by unphosphorylated PtsN, which interacts with the CorC domains with high affinity and allosterically augments inhibitory interactions of CorC with transport helices of YcgO. Inhibition is relieved leading to constitutive activation, upon disrupting the CorC-transport conduit interface. This study illuminates the structural basis of K^+^ efflux mediated through regulation of a K^+^/H^+^ antiporter in *E. coli* and related prokaryotes via a metabolic network involving a regulatory phosphorelay.

## Introduction

Ionic fluxes across cellular membranes are vital to sustain fundamental physiological processes, such as ionic homeostasis, osmotic adaptation, pH tolerance and maintenance of membrane potential^1,2^. Cation proton antiporters (CPAs) play a vital role in maintaining K^+^ and Na^+^ levels across cellular membranes^3,4^. Peter Mitchell proposed the existence of electroneutral K^+^/H^+^ and Na^+^/H^+^ antiporters in cellular membranes to overcome the deleterious effects of overaccumulation of monovalent cations within the cytosol due to the negative membrane potential imposed by the pH gradient formed during oxidative phosphorylation^5^. While multiple Na^+^/H^+^ antiporters have been structurally characterized, very few K^+^/H^+^ antiporters are structurally and mechanistically understood^6–8^. A recent structural study of KefC, a CPA2 K^+^/H^+^ antiporter, that responds to electrophiles through interactions with electrophile generated glutathione-S-conjugates, provides insights into K^+^-specific antiport^8^. Unlike Na^+^/H^+^ antiporter structures that primarily comprise a membrane embedded transporter, some K^+^-specific antiporters possess additional cytosolic domains to regulate their function typified by the homodimeric KefC, that has cytosolic RCK domains linked to the transporter^7–10^. A CPA1 K^+^/H^+^ antiporter from *Vibrio cholerae,* VcNhaP2 related to YcgO, from *E. coli* was functionally characterized to be involved in electroneutral K^+^ efflux^11^.

We earlier uncovered the interplay between a three protein phosphorelay comprising PtsP-PtsO-PtsN and K^+^ efflux mediated by the integral membrane protein YcgO. YcgO, an integral membrane protein displays sequence homology to Vc-NhaP2, an electroneutral K^+^-specific CPA^11^. Studies by us and others demonstrated that in wild-type *E. coli* K^+^ efflux by YcgO is inhibited by the unphosphorylated form of PtsN (PtsN_UnP_), the terminal phospho-acceptor protein of the PtsP-PtsO-PtsN phosphorelay^12–14^. Absence of PtsN_UnP_ led to the paradoxical phenomenon of YcgO mediated cellular K^+^ limitation in media containing high external K^+^ concentrations, that are more than sufficient for optimal growth^12,14^. Inhibition and/or inactivation of the high affinity K^+^ uptake system, KdpFABC, by high but not low external K^+^ is thought to be a factor in maintaining K^+^ limitation in the Δ*ptsN* mutant^12,15^. PtsN_UnP_ also interacts with KdpD/E two component system and enhances the expression of KdpFABC uptake system^16,17^. K^+^ limited growth of the Δ*ptsN* mutant could be overcome either by removal of YcgO or by overexpression of the Kup K^+^ uptake transporter^12^.

Additionally, we identified that PtsN_UnP_ exerts its inhibition of YcgO by interacting with its cytosolic CorC domain, an interaction attenuated by PtsN phosphorylation^13^. It is suggested that the K^+^/H^+^ antiport function of YcgO may represent adaptation to as yet unidentified stress conditions^12^. A recent study has shown that the phosphorylation states of PtsN can modulate resistance to the toxic electrophile methylglyoxal via YcgO mediated K^+^/H^+^ antiport^18^.

In this study, we elucidate the high-resolution cryoEM structures of the complex between YcgO and PtsN_UnP_ and quantify the interaction to reach nanomolar affinity. The interaction is in the midst of the cytosolic domains of YcgO that comprise an RCK domain^19^ linked to a CorC/HlyC domain (CorC)^20^. The CorC domain is beneath the helices of the elevator/transport domain forming additional interactions with the transmembrane helices of YcgO involved in gating. A phosphomimic version of PtsN fails to interact with YcgO and a structure of YcgO devoid of PtsN_UnP_ displays reduced order within the cytosolic domains. The transporter in both states is bound to a K^+^ ion in the ion-recognition site which remains occluded from solvent access from both the extra and intracellular environments. Disrupting the K^+^-binding site through an efflux deficient D162N substitution results in a deficiency in K^+^/H^+^ movement. Our study supports a model which postulates that constraint on the CorC domain mobility imposed by bound PtsN_UnP_ engenders allosteric inhibitory interactions of this domain with the transport helices of YcgO. These studies establish the mechanism underlying the metabolic control of K^+^ antiport in YcgO by a regulatory phosphorelay.

## Results

### CryoEM structure of the YcgO dimer

YcgO is closely related to the CPA1 class of K^+^/H^+^ exchangers ubiquitously present in numerous Gram-negative bacterial species (Extended Data Figures 1, 2a). Functional analyses of a *Vibrio cholerae* homologue Vc-NhaP2, revealed this transporter class to be a K^+^/H^+^ electroneutral antiporter bearing the CPA1 motif for interacting with K^+^ ions and involved in electroneutral transport (Extended Data Figure 1). A C-terminally 3x FLAG hexahistidine tagged YcgO (YcgO_FH_) used for all cryoEM analyses and referred to as YcgO in the context of these and other biochemical studies, was expressed and purified from the *E. coli* strain C41 and extracted from the inner membrane fraction using lauryl maltose neopentyl glycol (LMNG) (Extended Data Fig 3). YcgO was enriched using affinity chromatography and purified to homogeneity using size exclusion chromatography (Extended Data Fig. 3a, b). Single particle cryoEM data were collected using a 300 keV Titan Krios with a biocontinuum K3 detector through which the structures of YcgO and YcgO bound to PtsN_UnP_ were reconstructed and refined to global resolutions of 3.4 Å and 3.2 Å, respectively (Extended Data Figures 4, 5).

The cryoEM structure of YcgO revealed a homodimeric organization typical of a CPA transporter with each protomer bearing 13 transmembrane (TM) helices (Fig. 1). Due to weak density observed for the cytosolic domains, the model was built primarily for the TM region of the transporter from residues 1-396. The N-terminus of TM1 faces the periplasm and is involved in mediating interactions within the dimer (Fig. 1a, b). Helices 1, 3, 8 and 10 form the dimeric interface with a hydrophobic cavity amidst these helix densities with phospholipid densities into which we modeled two phospholipid molecules of palmitoyl oleoyl phosphatidyl ethanolamine (POPE) located primarily in the outer leaflet of the membrane (Fig. 1b, c; Extended Data Figure 6; Supplementary Figure 1). Dimer interactions between TM helices constitutes 1820 Å^2^ of interfacial area for each protomer that is near 10% of the total surface area for each protomer within the plane of the membrane. Ordered cytosolic domains, particularly the RCK domains, participate in enhancing the dimerization interface (described later). Each of the modeled phospholipid has an interfacial area of 620 Å^2^ with the two YcgO protomers. Phospholipids are observed to stabilize the interfaces of dimeric CPAs like KefC, KimA and PaNhaP^7,8,21^. Since extraneous lipids were not added during extraction, it is assumed that the phospholipids observed in the map are native lipids extracted alongside the dimeric YcgO. The topology of TM helices in YcgO can be clearly divided into core/dimerization domain/helices (TMs 1-3; 7-10) and transport domain/helices (TMs 4-6; 11-13) (Fig. 1c, d). The dimerization domain is expected to act as a scaffold allowing helices in the transport domain to undergo alternating-access as an elevator to complete the transport cycle. The TMs 5 and 12 in YcgO are unstructured within the bilayer to form an “X”-shaped configuration to allow ion coordination and transport. The motif formed by this disorder within TM helices is a conserved structural feature among all members of the CPA family and ion-specificity is controlled to changes in the motifs that surround the ion-binding site^22^. The activity of YcgO overexpressed in *E. coli* membranes was evaluated using ACMA (9-amino-6-Chloro-2-methoxyacridine) fluorescence as a reporter for pH gradients across the vesicle membranes. The assay employed an antiporter deficient strain of *E. coli* (GJ22224) lacking YcgO, KefC, YbaL KefB, NhaA, ChaA and PtsN (Fig. 1e). This allowed the analyses of activity of a functional overexpressed C-terminally 3x FLAG tagged YcgO,^13^ (YcgO_WT_) in everted vesicles prepared from GJ22224, using ATP to quench ACMA fluorescence, signaling the formation of pH gradient^13^. Consistent with an earlier report^13^, addition of 5 mM KCl induced a dequenching of ACMA fluorescence in vesicles with YcgO_WT_ indicating formation of a proton flux in response to K^+^ addition. Addition of 5 mM Rb^+^, a congener of K^+^,^23^ also induced a clear and specific proton flux across the membrane (Fig. 1f, Supplementary Figure 2a). Comparison of the TM region of YcgO with other CPA transporters with known structures like KefC, MjNhaP, EcNhaA and TtNapA reveal that YcgO is closest in structural overlap with MjNhaP structure at neutral pH compared to structures of other CPA members (Extended Data Figure 7).

**Figure 1:**
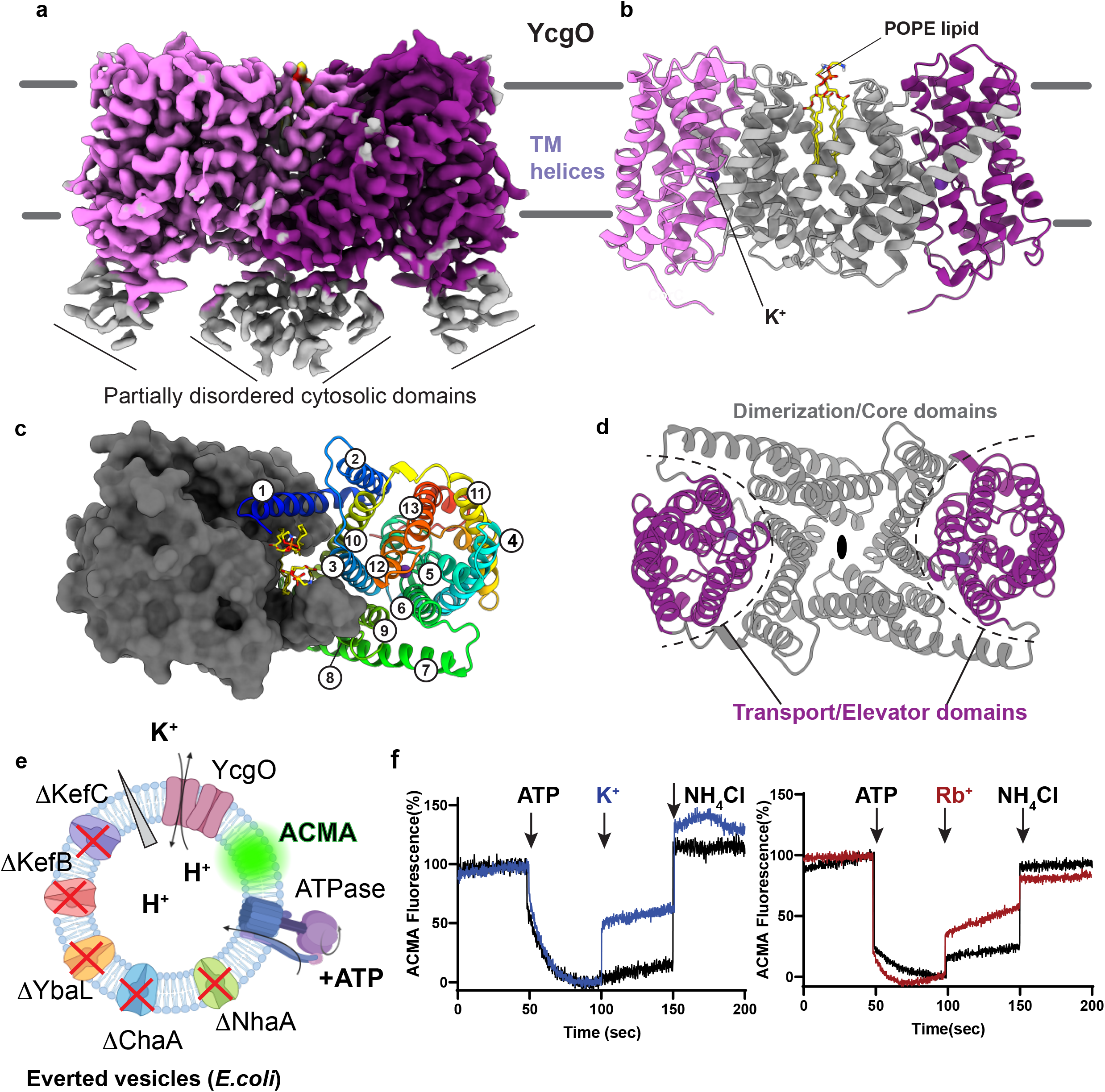
CryoEM structure of YcgO. **a,** Density of YcgO resolved to 3.4 Å displaying TM helices but partially disordered cytosolic domains (contoured at 0.16 σ). The two protomers of the dimer are colored in pink and purple and disordered cytosolic domains are colored in gray. **b**, Modeled structure of YcgO TM helices display a dimer with phospholipids (POPE, yellow) in the interface. The TM helices involved in dimerisation are colored in gray and helical bundles involved in elevator mechanism are colored in pink and purple. **c**, Top-down view of YcgO dimer displaying one protomer as a surface (dark gray) and helices in rainbow colors with TM1 (blue) to TM13 (red). Interfacial lipids can be observed **d,** Top-down view depicting two-fold axis and demarcation between dimerisation and transport helix bundles. **e,** Schematic of the everted vesicle experiment performed to demonstrate proton flux across vesicle membranes upon addition of alkali metal ions. Proton gradient is represented in a gray arrowhead. Other secondary active transporters knocked out of the *E. coli* strain GJ22224, are represented with red crosses. Schematic was prepared using Biorender. **f**, Everted vesicle K^+^ efflux demonstrated with membrane vesicles from the strain GJ22224 that overexpresses a functional, C-terminally 3x-FLAG tagged YcgO and vesicles prepared from the same strain bearing empty vector. ACMA fluorescence was used to monitor changes to ATP induced pH gradient (50 s) and 5 mM K^+^-induced dequenching (100 s) of fluorescence indicating H^+^-flux across the membrane followed by quenching of pH gradient with the use of NH_4_Cl (150 s). Rb^+^ was also tested for specific ability to dequench ACMA fluorescence. Empty vesicles in both cases respond with minimal dequenching to K^+^ and Rb^+^ addition. One of 2 technical repeats is represented from a set of four biological repeats (n=4).

### CryoEM structure of YcgO complexed with unphosphorylated PtsN

We assessed the *in vivo* functionality of YcgO_FH_ used for all cryoEM analyses. For this purpose, the gene encoding YcgO_FH_ was placed at its native genomic location (described in supplementary methods). In the wild-type state a strain expressing a chromosomally encoded untagged YcgO grows equally on low (1 mM, K_1_) and high (115 mM, K_115_) K^+^ containing, glucose supplemented, synthetic media. Absence of PtsN leads to K^+^ limited growth inhibition in K_115_ medium owing to YcgO mediated K^+^ efflux, that is alleviated either by removal of YcgO or by overexpression of the Kup K^+^ uptake transporter^12^ (Supplementary Figure 3). The gene encoding YcgO_FH_ displayed all the above phenotypes attributed to the untagged YcgO, indicating that the appended bipartite tag does not alter function of the YcgO construct used or the cryoEM studies (Fig. 2a, Supplementary Figure 3). The functionality of C-terminally hexahistidine tagged PtsN is described earlier^12,13^. For the purposes of cryoEM and biochemical studies unphosphorylated C-terminally hexahistidine tagged PtsN is also referred to as PtsN_UnP_. Purified YcgO and PtsN_UnP_ display interactions when mixed resulting in a leftward shift of the YcgO peak in fluorescence-detection size exclusion chromatography (FSEC) observed using tyrosine fluorescence (λ_ex_=275 nm; λ_em_=305 nm, Fig. 2b). The 2D classes of YcgO complex revealed the extra density corresponding to bound PtsN_UnP_ and structure refinement yielded a map at a resolution of 3.2 Å (Fig. 2c, d; Extended Data Fig. 5; Supplementary Figure 4). PtsN_UnP_ was added prior to grid freezing at a ratio of 1:5 to ensure saturation of the YcgO sample with bound PtsN_UnP_. The structure of the complex yielded a resolution of 3.2 Å with clear densities for the cytosolic RCK domains, CorC domains and for the bound PtsN_UnP_. Side chains could be modeled and refined without any bias using the AlphaFold2 model of YcgO and the crystal structure of PtsN (Extended Data Table 1). The TM helices in the YcgO-PtsN_UnP_ complex are organized very similar to the TM regions of the PtsN_UnP_-free YcgO structure. The densities for phospholipid molecules were visible in the complex structure and POPE was modeled into the visible densities (Fig. 2d, e). The dimerization interface is accentuated in the presence of RCK domains in the cytosol and the interfacial area increases to ∼2920 Å in the full length YcgO dimer. The complex between YcgO-PtsN_UnP_ highlights the multidomain organization of YcgO and consequently the extensive interfaces it has in the complex with PtsN_UnP_. Close knit homodimeric interactions are observed amongst YcgO TM helices and RCK domains. PtsN_UnP_ interacts with YcgO in the wedge between RCK and CorC domains with primary interactions with the CorC domain (Fig. 2e, f, Extended Data Fig. 6). Both YcgO and PtsN_UnP_ have several closely related orthologues in Enterobacteriaceae. It is therefore likely that the role of YcgO mediated K^+^/H^+^ antiport is a highly conserved phenomenon across numerous bacterial species (Extended Data Figure 2).

**Figure 2:**
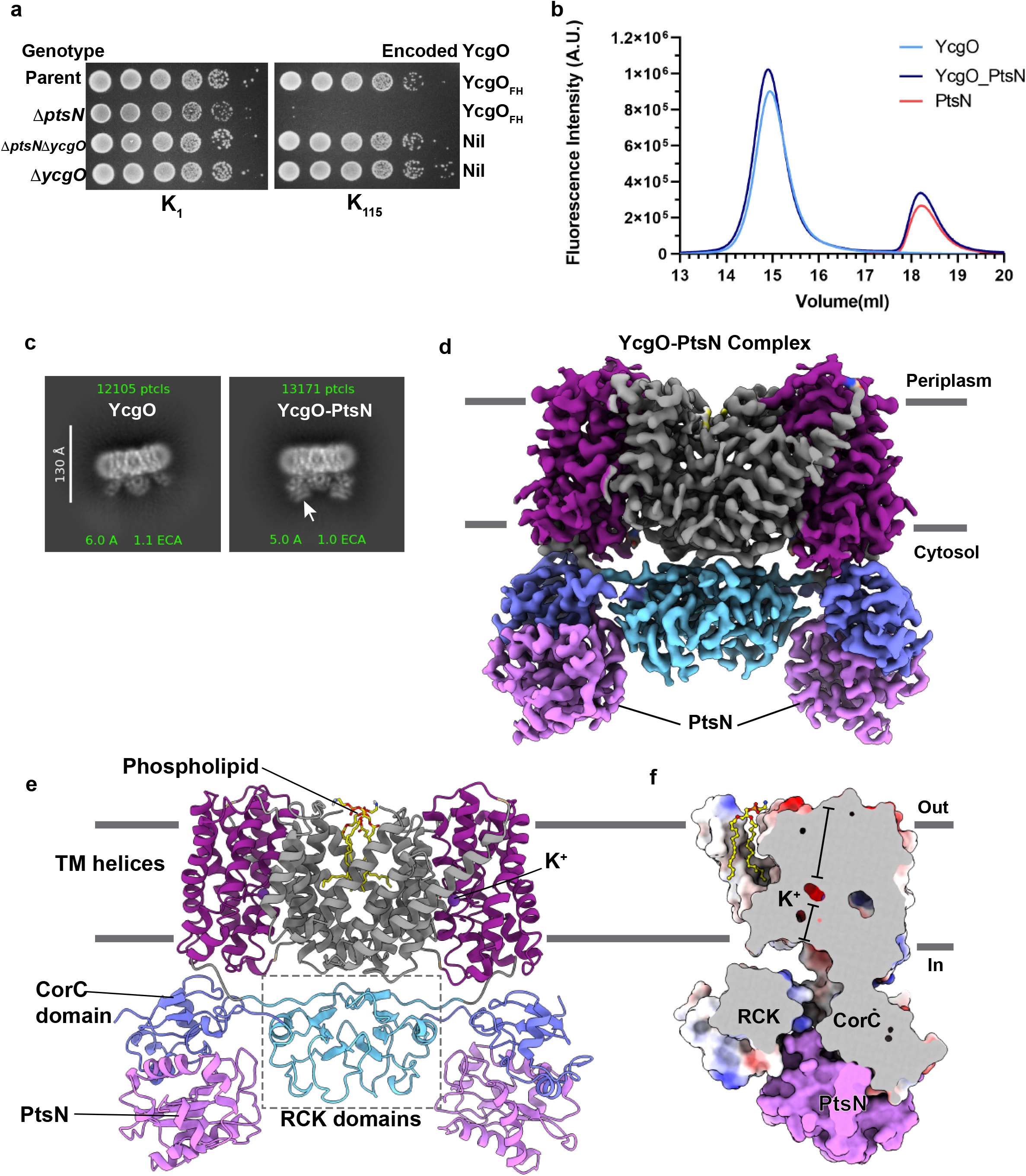
CryoEM Structure of the YcgO-PtsN complex. **a**, Functionality of the C-terminally 3x-FLAG and 6x-histidine tagged YcgO (YcgO_FH_) construct used for CryoEM structure determination. Cultures of the wild-type (Parent) expressing chromosomally encoded YcgO_FH_, its Δ*ptsN* derivative (Δ*ptsN*) and derivatives of the parent lacking YcgO alone (Δ*ycgO*) or in combination with the Δ*ptsN* mutation (Δ*ptsN* Δ*ycgO*), were ten-fold serially diluted and spotted on the surface of minimal glucose agar plates with K^+^ concentrations of 1 mM (K_1_) and 115 mM (K_115_), respectively. Strains employed are GJ22301 (Parent), GJ22302 (Δ*ptsN*), GJ22206 (Δ*ptsN* Δ*ycgO*) and GJ19262 (Δ*ycgO*) and are listed in the supplementary information section. One of two independent biological repeats is represented in the figure. **b,** Purified PtsN_UnP_ can shift the YcgO profile to a higher hydrodynamic radius as observed using tyrosine fluorescence (λ_Ex_=275 nm, λ_Em_ =305 nm). **c,** Reference-free 2D class comparison between YcgO and YcgO-PtsN_UnP_ complex reveals extra density that corresponds to bound PtsN_UnP_, indicated in the figure as PtsN. **d,** Refined map of YcgO dimer and PtsN_UnP_ interacting with its cytosolic domains, resolved to 3.2 Å. Map is colored according to the domains indicated. **e**, The refined model built into the density reveals a K^+^-bound occluded transporter with cytosolic RCK (cyan) and CorC domains (pale blue) linked through a disordered linker. PtsN_UnP_ (lavender) sits between and interacts with both the RCK and CorC domains. CorC domains in the cytosol closely interact with the transport helices of YcgO. **f,** A sagittal section of the YcgO protomer reveals bound K^+^-buried deep within the TM helices in a site occluded from solvent access from both extracellular and intracellular compartments.

Interestingly, we could observe clear density for the potassium ion in the middle of membrane plane in both PtsN_UnP_-bound and free structures of YcgO (Fig. 3). The bound ion was occluded from solvent access from both the extracellular and intracellular faces of the transporter. Occluded states of CPA transporters have not been reported earlier, to the best of our knowledge. We therefore compared the inward-open state of KefC with YcgO and observed subtle variations in helix positions that occlude the ion from solvent access. The dimerization helices of TMs 3 and 10 tilt towards the cytosolic vestibule. The disordered region of TM12 that harbors the K^+^-binding site also displays a displacement of a 5.0 Å relative to KefC TM12 linker despite K^+^ ion in a nearly overlapping position. TM13 in YcgO displays a tilt from 381-390 that allows the lowest part of this helix to turn towards the vestibule. This helix is connected to the C-terminal linker to the cytosolic RCK and CorC domain and could allow the allosteric opening and closing of the vestibule in response to dynamics within the cytosolic domains. These differences in the organization of TM helices between KefC and YcgO effectively contribute to the formation of an occluded state for K^+^ binding within YcgO (Extended Data Fig. 7e).

**Figure 3:**
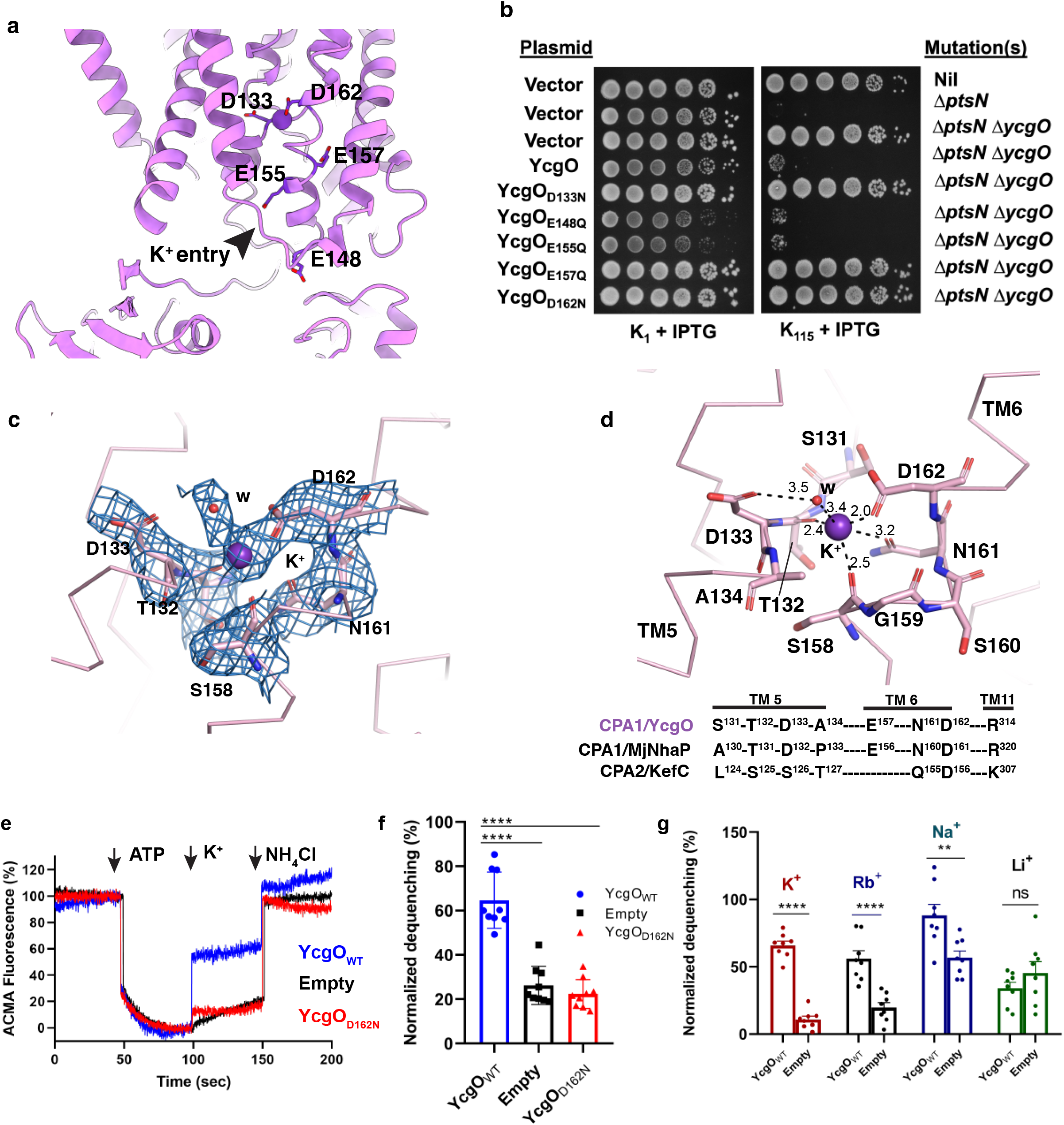
Role of acidic residues in ion coordination and transport in YcgO. **a,** Acidic residues lining the K^+^ entry vestibule from the cytosolic side of the transporter which leads to the primary ion-binding site. **b,** *In vivo* functionality of YcgO bearing substitutions of the indicated acidic residues. Cultures of the parent (Nil), its Δ*ptsN* derivative and the Δ*ptsN* Δ*ycgO* derivative bearing either the empty vector (pTrc99A) or pTrc99A expressing YcgO and its indicated substitution derivatives, were spotted on K_1_ and K_115_ glucose minimal agar plates containing 10 µM IPTG. All YcgO proteins are expressed from the plasmid borne P_trc_ promoter and bear a 3x FLAG epitope tag at their N-termini. Strains used are GJ17829 (Nil), GJ17831 (Δ*ptsN*) and GJ22206 (Δ*ptsN* Δ*ycgO*). Absence of growth in mutants indicates active state of YcgO and normal growth across dilutions indicates inactive transport. **c,** Density of the bound K^+^ and its coordinating residues and water. **d,** K^+^ ion coordinated at the interface of TMs 5 and 6 and coordinated through a motif that is primarily built in the midst of TMs 5 and 6. Sequence alignment snippets of ion coordinating residues observed in other CPA members. **e,** Everted vesicles derived from the strain GJ22224 displaying the K^+^ intake in YcgO using ACMA fluorescence changes (Ex= 409 nm; Em= 474 nm), indicate lack of K^+^ /H^+^ antiport activity in YcgO_D162N_ mutant as seen through everted vesicle-based K^+^ efflux assay. **f,** Quantification of everted vesicles-based K^+^ efflux activity across multiple replicates for YcgO_WT_ and YcgO_D162N_. 10 mM KCl was added to induce H^+^ flux in this experiment. YcgO_WT_ displays substantial dequenching of ACMA fluorescence and YcgO_D162N_ is its D162N substitution derivative is inactive. Error bars represent S.E.M and data points indicate each of the repeats done for this experiment with three technical repeats done for every biological repeat (n=3). Comparison dequenching observed with K^+^ with vesicles containing YcgO_WT_, empty control and YcgO_D162N_ are indicated by four stars indicating a *p*-values of less than 0.0001 using ordinary one-way ANOVA. **g**, Effect of alkali ions tested using K^+^, Rb^+^, Na^+^ and Li^+^ were also checked by adding 5 mM of each ion to everted vesicles done using 4 biological repeats (n=4) each performed in duplicate. All points were used to plot the histogram. Error bars represent S.E.M and unpaired t-test indicated the *p* values of <0.0001 for K^+^, <0.0001 for Rb^+^, 0.0051 for Na^+^ and 0.248 for Li^+^ when compared with empty vesicles.

In the PtsN_UnP_ free state, the CorC and RCK domain densities were partially visible due to the likely disorder within the PtsN_UnP_-free YcgO structure that did not allow appropriate modeling of the atomic coordinates within these densities. However, both the RCK and CorC domains were observed very clearly in the PtsN_UnP_ bound complex likely due to the constraint induced by the binding of PtsN_UnP_. The RCK (396-483) and CorC (490-575) domains, could be modeled without any breaks in the main chain. The RCK domain is linked to TM13 with long unstructured linker which could be modeled into the visible density. The RCK domain of YcgO has similarities with RCK domains found in other bacterial K^+^ channels where they regulate the opening and closure of K^+^ channels in both prokaryotic and eukaryotic K^+^ channels. An overlap of an c-di-AMP bound RCK domain crystal structure from a *S. aureus* CPA transporter (SaCpaA) (uniport ID. Q2FZQ4) onto the RCK domain dimer of YcgO yields an overlap of the individual domains with an RMSD of 1.8 Å for 68 Cα atoms^24^. The c-di-AMP molecule bound within the RCK domain dimer of SaCpaA reveals a conservation of the binding site organization within the two RCK domain of YcgO. However, the binding site has some critical substitution like T456 (RCK_YcgO) in place of H184 (RCK_SaCpaA) and R442 (RCK_YcgO) in place of I170 (RCK_SaCpaA) that would compromise the ability of the RCK domains in YcgO to interact with secondary metabolites like c-di-AMP. The positions of the binding site in RCK domains in YcgO is substantially separated by about 25 Å (R433 Cα-Ca distance), whereas the separation within the SaCpaA RCK domains with bound c-di-AMP is about 12.6 Å (Extended Data Fig 8a). These features likely impair the abilitiy of nucleotides to interact with the YcgO RCK domains. The variations in the RCK domain of YcgO that would compromise its ability to bind to c-di-AMP in comparison to the SaCpaA RCK domain, can be rationalized on the basis that *E. coli* does not synthesize c-di-AMP. Further an overlap of the YcgO RCK domains with the KefC RCK domain displays a substantial difference in the organization of the two domains. The RCK domain in KefC is quite extensive and harbors glutathione and nucleotide binding sites but has very minimal overlap with the RCK domain of YcgO (Extended Data Figure 8b).

The CorC domain is connected to the RCK domain with another unstructured linker. The residues from 490 to 575 constitute the CorC domain that has structural similarities to the C-terminal CorC/HlyC domain from the Mg^2+^ transporter CorC C-terminal domain^20^. The domain is also observed in multiple proteins like hemolysin C (Extended Data Fig. 8). The domain was not attributed any obvious function prior to this study. However, in YcgO the domain is primarily involved in interactions with PtsN_UnP_ and can allosterically affect the function of YcgO (described later).

### Ion coordination within YcgO

Protomers in both structures of YcgO reported in this study retain a clear density for a bound monovalent ion in the ion binding site of YcgO. A K^+^ ion was modeled into this density within the dimer as the major salt in the buffer being KCl during structure determination of YcgO. The ion is surrounded by acidic residues and additional acidic residues are observed along the vestibule leading up to the ion-binding site (Fig. 3a). We evaluated the roles of acidic residues involved in ion coordination and those lining the vestibule for effect on YcgO function *in vivo*. Phenotypic studies employed here tested for the ability of substitution bearing derivatives of a functional N-terminally 3x FLAG tagged YcgO,^13^ to complement the Δ*ycgO* mutation in the absence of PtsN. In this background, a functional YcgO leads to growth inhibition in K_115_ medium^13^ (Fig. 3b). Antiporters involved in cationic substrate transport tend to have acidic residues in the vestibule for protonation-driven conformational transitions^25,26^. We observe that cells carrying YcgO substitutions E148Q, E155Q, that line the vestibule, do not affect the activity YcgO (Fig. 3b). On the other hand, isosteric modifications in the conserved CPA1 motif at residues D133, D162 and E157, to N, N and Q respectively, lead to growth in K_115_ medium, indicative of inactivation of YcgO (Fig. 3b). This indicates the importance of the role of conserved acidic residue in this family of CPA1 transporters. D162 is directly involved in K^+^ coordination along with the N161 side chain and the carbonyl groups of T132 and S158 (Fig. 3c, d). D133 is involved in indirect coordination by interacting with a water molecule that coordinates K^+^ ion. Unlike KefC, we do not observe a complete dehydration of the bound ion in YcgO. The mean coordination radius of K^+^ ions is 2.8 Å which is very similar to K^+^ coordination found across multiple ion-bound structures^27^. Although E155 is not part of the coordination, it has a vital structural role in forming a conserved salt bridge with R314 in TM11 that stabilizes the local structure around this region (Fig. 3b, d). We performed everted vesicle assays to biochemically demonstrate that D162N inactivates YcgO and is incapable of dequenching the ACMA fluorescence indicating a lack of proton flux across YcgO D162N (Fig. 3e, f). YcgO_D162N_ was expressed at levels comparable to wild-type YcgO (YcgO_WT_, Supplementary Figure 2b). We further evaluated the ability of other monovalent cations for uptake activity using Rb^+^, Na^+^ and Li^+^ as candidate ions to observe for proton flux in everted vesicles. The K^+^ congener Rb^+^ displays a clear ability to dequench ACMA fluorescence indicating its ability as a substitute for K^+^ to induce proton flux in everted vesicles carrying YcgO. Na^+^ addition induced a high dequenching but the empty vesicles also had a substantially high ability to dequench fluorescence. However, a smaller ion like Li^+^ does not yield any difference in dequenching between YcgO_WT_ and empty vesicles suggesting absence of YcgO induced proton flux. This Na^+^ effect is likely because of background Na^+^ exchange occurring in the bacterial vesicles due to the activity of NhaB that could not be eliminated as a NhaB knockout in the strain GJ22224 led to lethality ^13^. The presence of S-T-D-A (YcgO) or L-S-S-T (KefC) in the CPA motif of K^+^-specific transporters as opposed to P-T-D-P motif predominantly observed in Na^+^-specific transporters is suggested to facilitate K^+^-specificity within CPA transporters^4^ (Fig. 3d). The motif definition may not be entirely foolproof in terms of ion specificity as we observe marginal levels of Na^+^ transport in YcgO and also reported for its homologue Vc-NhaP2^11^. However *in vivo*, the predominant ion that would be transported is likely to be K^+^ due to its excess presence within the cytosol.

### PtsN_UnP_ primarily interacts with the CorC domain

PtsN_UnP_ wedges into the space between the cytosolic RCK and the CorC domains in the cytosolic face of the transporter (Fig. 2f). The interfacial area between PtsN and CorC-RCK domain junction is 1190 Å^2^ with a bulk of the interactions occurring between CorC domain and PtsN_UnP_. The interaction stabilizes the two cytosolic domains. The CorC domain has close structural overlaps with the CorC domain from Nitrosomonas and Neisseria species (Extended Data Figure 8c, d). It also overlaps closely with a similar domain in hemolysin C and is also present as a C-terminal domain in the Mg^2+^ transporter CorC^20^ (Extended Data Figure 8e). Despite its presence in numerous proteins the exact role of this domain remains uncharacterized in these species. Molecular simulations of the YcgO model with the cytosolic domains were performed in the presence and absence of PtsN_UnP_. The simulations clearly revealed extensive flexibility in the CorC domain whereas the TM helices and the RCK domains remain relatively stable. A similar effect is also observed in the density of RCK and CorC domains of YcgO in the EM maps. RCK domain density is partially structured whereas CorC domain displays substantial displacements in the PtsN_UnP_ free state. In the presence of the bound PtsN_UnP_ the cytosolic domains retain minimal flexibility as observed in the cryoEM structure of YcgO-PtsN_UnP_ complex (Extended Data Figure 9a, b). The interaction of PtsN_UnP_ also seems to minimize solvent access to the cytosolic vestibule as quantified using simulations. The PtsN_UnP_ free YcgO displays greater solvent access over the simulation time frame within YcgO whereas PtsN_UnP_ bound YcgO has minimal solvent access during the same duration of the simulation (Extended Data Figure 10a). We made substitution in a few residues that line this vestibule at S139A (TM5), D272A and W276A (TM10). However, none of these substitutions disabled YcgO function (Extended Data Figure 10b) highlighting that these residues do not affect the ability of monovalent ions to diffuse and interact with the binding site within YcgO.

The ability to reduce disorder within the cytosolic domains through interaction with PtsN_UnP_ reflects in the interaction affinity of YcgO with PtsN_UnP_ that we tested using isothermal titration calorimetry (ITC). Both YcgO and PtsN_UnP_ were purified in the same buffer containing 0.1 mM LMNG to prevent heat changes due to dilution effects. The titration of PtsN_UnP_ into the cell containing YcgO yields an endothermic reaction causing a positive differential power (dP) that yields a titration curve whose analyses reveals an affinity of 188 nM (Fig. 4b, d). The enthalpy (ΔH) in this reaction is 8 kcal/mol whereas the TΔS value is 17.0 (kcal/mol) with a difference in free energy (ΔG) of binding calculated at −8.9 kcal/mol (Fig. 4b, d). The values indicate an entropically driven binding reaction that is a likely outcome of PtsN_UnP_ interaction with RCK and CorC domains displacing ordered water molecules around these domains prior to PtsN_UnP_ interaction^28^. PtsN_UnP_ interacts with the CorC domains of YcgO using helices α2, α3 and strands β1 and β3 (Fig. 4a). The long loop that connects β2 and β3 strands forms the minimal interface of PtsN_UnP_ with RCK domain (Fig. 4a). The β1 strand harbors a histidine residue (H73) that forms close interactions with Y514 and F499. Interestingly disruption of this interface in our earlier study through F499Y substitution yielded a constitutively active YcgO incapable of interacting with PtsN_UnP_ (Fig. 4a, inset)^13^. H73 in the interface of the PtsN and CorC domains is known to be the only site of phosphorylation in PtsN^29,30^. A phospho-mimic mutant of PtsN, H73E, was made to test its interaction propensity with YcgO. PtsN_H73E_ failed to interact with YcgO under identical conditions used for the ITC experiment yielding no quantifiable heats to fit a binding isotherm. These observations thus, reveal the importance of the PtsN_UnP_-CorC interfacial interactions in the control of K^+^ efflux in YcgO, which can be perturbed by phosphorylation of PtsN (Fig. 4c).

**Figure 4:**
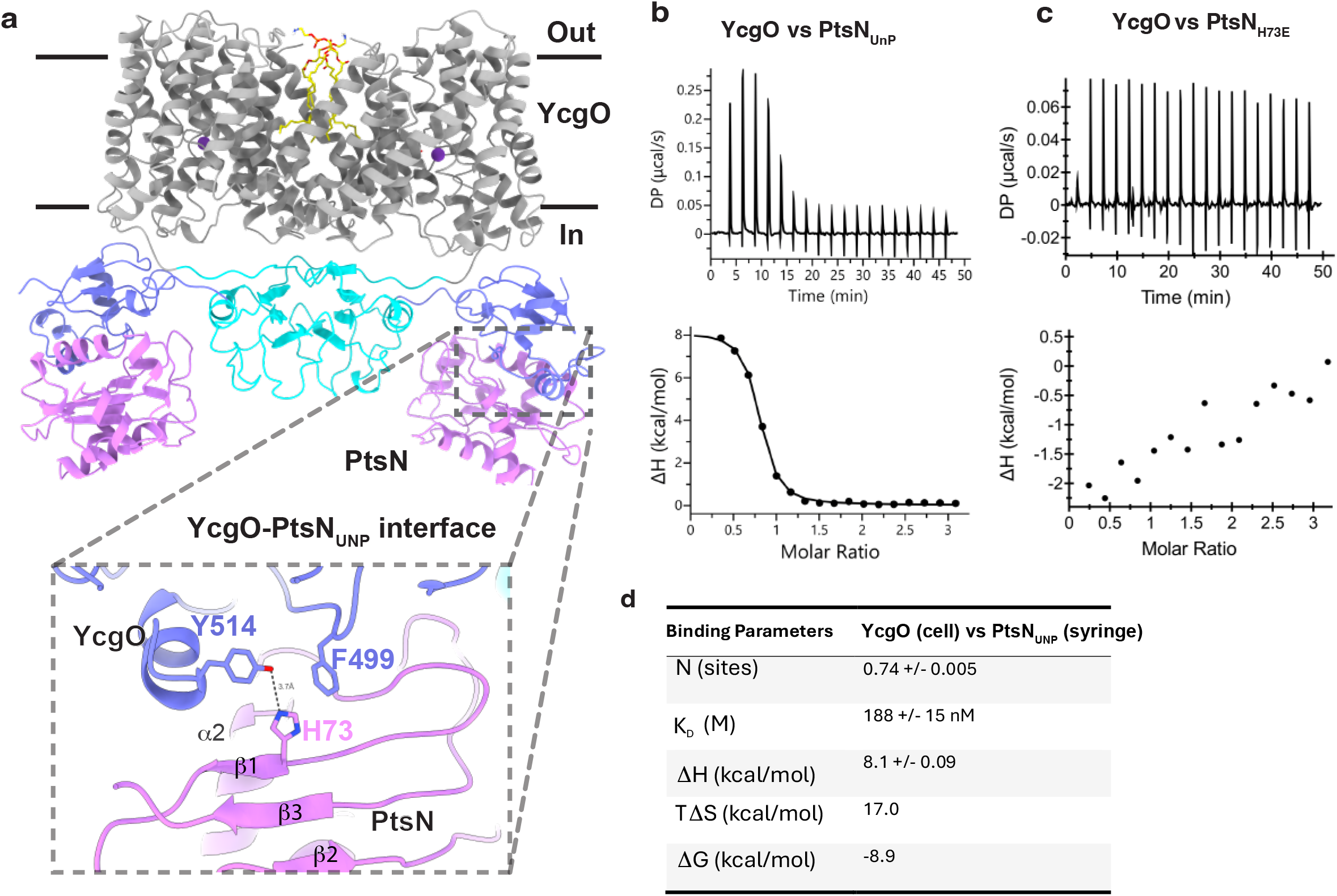
YcgO-PtsN interface disruption through phosphorylated PtsN interaction. **a,** YcgO-PtsN_UnP_ (PtsN) complex interface, with emphasis on interfacial residues in the CorC domain. PtsN H73’s position and its interaction with Y514 and F499 (F499Y is a site for constitutive activation) at the interface (inset) is depicted to convey the lack of space for accommodating a phosphate moiety in the event of H73 phosphorylation. **b,** Isothermal titration calorimetry reveals a dissociation constant of 188 nM for the YcgO-PtsN_UnP_ interaction. Heat of dilution was subtracted prior to analyses. One of two biological repeats is represented in the figure. **c,** A phospho-mimic substitution of H73E completely nullifies the YcgO-PtsN interaction. **d,** Table depicts binding parameters for the YcgO-PtsN titration from panel c for ITC data. The YcgO protein used is YcgO_FH_.

### CorC domain exerts allosteric control on YcgO transport helices

The CorC domain that has the predominant interactions with PtsN_UnP_ also has close-knit interactions with the helices of YcgO that are involved in the elevator-based movement within YcgO (Fig. 5a). The loop between helix α3 and strand β1within the CorC domain is the primary site of interaction with the loops amidst transport helices connecting TMs 5 and 6 followed by TMs 11 and 12. These TMs are vital for harboring the K^+^ binding site and are involved in alternating-access to transport K^+^ from the cytosol to the periplasm. The interaction is supported through hydrophobic interactions but also an electrostatic interaction between R149 (cytosolic loop between TM5-TM6) and D545 (CorC) and E148 (TM6) and K559 (CorC) (Fig. 5a, inset1). We previously noted that the D545A substitution in the CorC domain of YcgO, led to constitutive activation of YcgO regardless of the presence of PtsN_UnP_ *in vivo*^13^. The D545A substitution would disrupt this salt bridge thereby decoupling CorC domain from the transport helices. The structure provides a rationale for this mutant phenotype as a detached CorC cannot effectively control YcgO inhibition modulated through PtsN_UnP_ interaction thereby activating antiport activity. As a further test of this notion, we prepared a chromosomal complementary R149A mutant that should display similar effect as YcgO_D545A_. Rather interestingly, chromosomal expression of YcgO_R149A_ yields a similar constitutive activation of YcgO, as does a previously reported YcgO_ΔCorC_ derivative that lacks the CorC domain^13^. Chromosomal expression of YcgO_R149A_, YcgO_D545A_, YcgO_ΔCorC_ but not YcgO in a *ptsN*^+^ background led to growth inhibition in K_115_ medium (Fig 5b). Inset 2 in figure 5 depicts an analogous abrogation of transporter interaction in absence of the CorC domain. Both YcgO_D545A_ and YcgO_ΔCorC_ fail to interact with PtsN ^13^.This provides vital insights into the crucial regulatory mechanism that PtsN_UnP_ interaction enforces on YcgO activity, and the vital and dual role played by CorC domain in control of YcgO activity.

**Figure 5:**
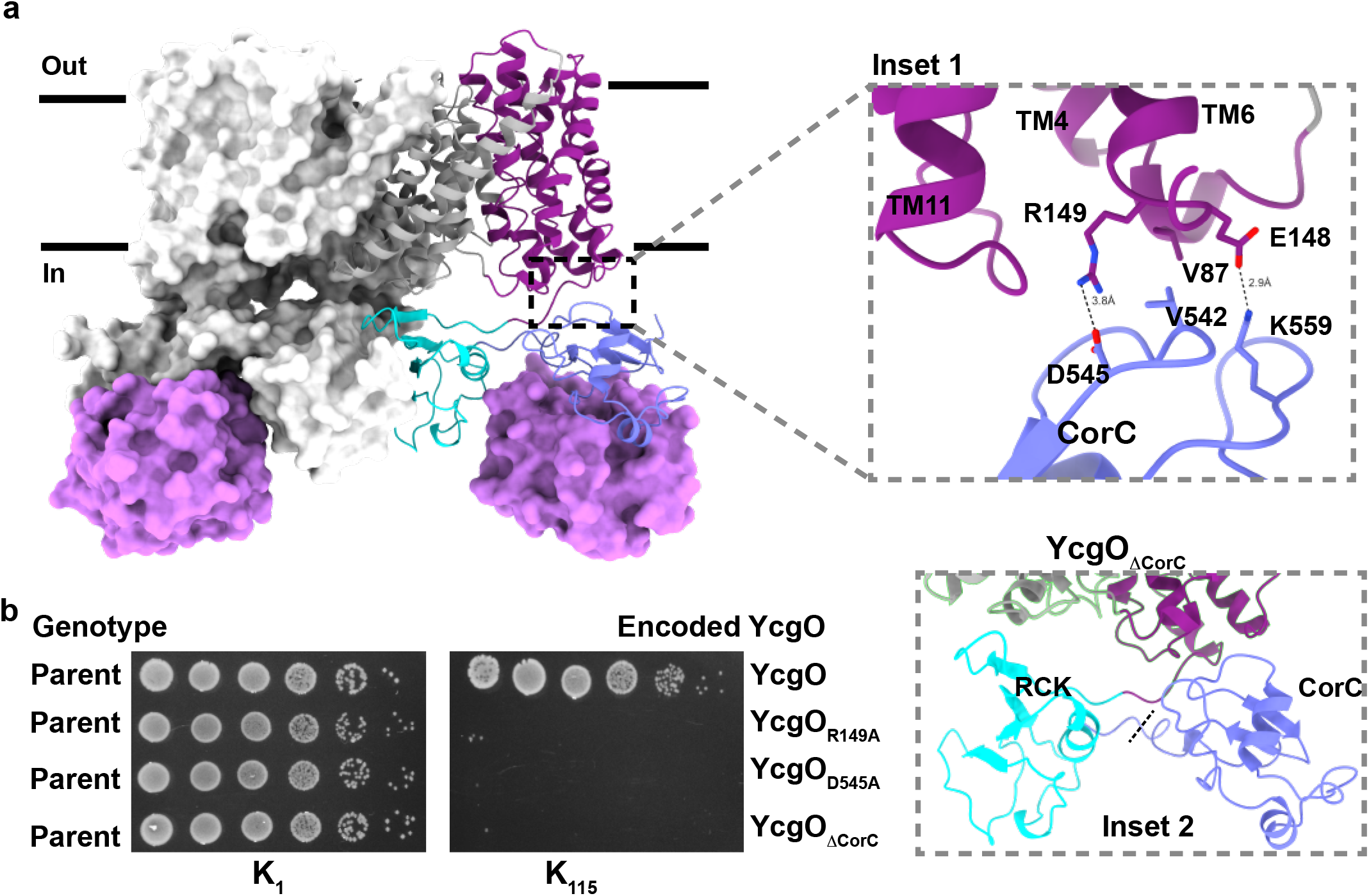
YcgO interdomain interactions between CorC and TM helices control transport activation. **a,** PtsN_UnP_ mediates YcgO inhibition by stabilizing inter-domain interaction within YcgO. A zoom-in inset 1 is shown for interfacial residues involved in interaction between elevator (purple) and CorC (pale blue) domains. H-bonds are demarcated with dashed lines. **b,** Constitutive activation of YcgO caused by the disruption of the D545: R149 salt bridge pair. Alanine substitutions at D545 and R149 yield constitutively active mutants. Cultures of wild type strain (Parent) chromosomally expressing a functional 3x FLAG tagged YcgO (YcgO) and its derivatives bearing either the indicated substitutions or a deletion removing the CorC domain (ΔCorC) were serially diluted and spotted on K_1_ and K_115_ agar plates. Inset 2 displays loss of CorC transporter domain interactions in YcgO_ΔCorC_ that lacks the CorC domain.

## Discussion

This study highlights the multidomain organization of YcgO through cryoEM structures and the roles that three domains of YcgO namely the transmembrane, RCK and the CorC domains of YcgO play in mediating K^+^ efflux. We have further reported the high-resolution structure of YcgO in complex with PtsN_UnP_. Mechanistically, the two structures reveal a K^+^ ion that is trapped in an occluded state, which has generally not been observed with other CPA transporters, whose structures are reported predominantly in either inward-open or outward open states^6–8,10^. Occlusion of the ion within the coordination site would render the K^+^ binding site of YcgO inaccessible to solvent or protons. The TM helices involved in ion transport in YcgO include TMs 4, 5, 6 and TMs 11, 12 and 13. TMs 5 and 12 form the ‘X’-shaped feature with discontinuous helices that form the coordination site along with residues within TM6. An analogous arrangement was reported for NhaA ^22^. The control of YcgO activity through PtsN_UnP_ interactions establishes a new mechanistic framework for K^+^efflux mediated by a member of the CPA protein family. While the transport is primarily facilitated by the transmembrane module of YcgO whose organization is similar to other CPA antiporters, the RCK and CorC domains uniquely influence this ability. Analyses on the YcgO-PtsN_UnP_ complex defines two functional and regulatory features of the CorC domain, the target of PtsN_UnP_ namely (i) the PtsN_UnP_-CorC interface and (ii) the PtsN_UnP_-CorC transporter interface. The former delineates the molecular basis for the differential abilities of phospho-PtsN versus PtsN_UnP_ to target the CorC domain. The H73 residue in PtsN forms the interface with the Y514 and F499 in YcgO. Phosphorylation of this site would disrupt this interface likely through steric clashes and greatly weaken the interaction of phospho-PtsN with YcgO causing unfettered K^+^/H^+^ antiport (Fig. 6). The inability of the F499Y substitution to interact with PtsN_UnP_,^13^ can thus be rationalized and also offers a physiological basis for why PtsN_UnP_ has an overwhelmingly high affinity for the CorC domain as opposed to its phospho-counterpart. *In vivo* in *E. coli*, PtsN_UnP_ is the minority species, phospho-PtsN is the major species^14,31^. It is suggested that a metabolic signal enhancing PtsN phosphorylation, would entail conversion of a minority fraction of PtsN_UnP_, to phospho-PtsN, leading to rapid adaptive K^+^ efflux^13^. The second regulatory effect of the PtsN_UnP_ CorC interaction is more intriguing and our studies show that the molecular outcome involves salt bridge mediated inhibition of the transport helices of YcgO, by the CorC domain, upon engagement with PtsN_UnP_. A notion supported by (i) the observation that perturbing this salt bridge leads to constitutive activation of YcgO and (ii) molecular simulations.

**Figure 6:**
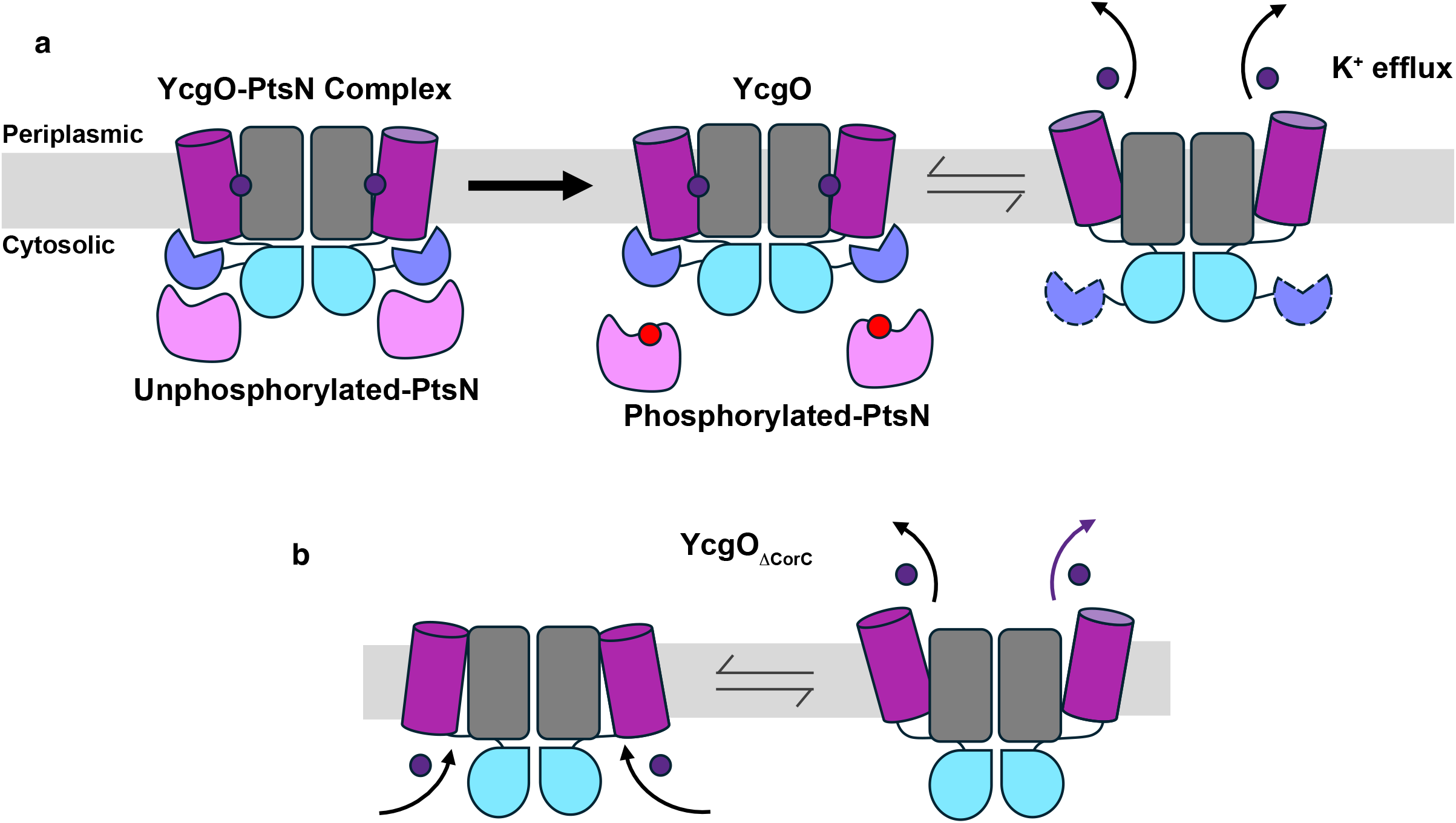
Schematic depicting regulation of YcgO mediated K^+^ efflux by unphosphorylated-PtsN. **a,** Unphosphorylated-PtsN interacts with high affinity largely with the CorC domain of YcgO. The unphosphorylated-PtsN bound CorC interacts with the transporter domain to keep the efflux process inhibited. Phosphorylation at H73 in PtsN through the PtsP-PtsO phosphorelay releases PtsN from YcgO resulting in the activation of K^+^-efflux. **b,** Altering the interface between CorC domains with the transport domain of YcgO through the ΔCorC deletion renders YcgO constitutively active.

The regulatory impact of the RCK domain in YcgO function is more enigmatic. In the classical examples, RCK domains are known to play a direct role in the cytosolic gating of K^+^ channels/transporters, in concert with their cognate small molecule regulatory ligands, whether they are tethered to or interact with the transporter domain as soluble proteins^32–34^. Whereas we could assign densities to bound POPE (Fig. 1, 2), we did not observe density for bound ATP at RCK, despite adding ATP in solution for the PtsN_UnP_ free YcgO. This does not preclude the existence of other endogenous ligands or even K^+^, to interact with the YcgO RCK domains to modulate YcgO function. Such regulation could have implication for K^+^ efflux as RCK is connected directly with the TM13 that could influence solvent accessibility at the cytosolic face of YcgO. We would like to consider another possibility. In the case of YcgO, a direct interaction of its RCK domain with the transporter module is less likely, in comparison to the related CPA KefC, where tethered RCK domains in conjunction with bound cognate ligands directly engage with the transporter module^8^. Furthermore, we noted that even in the absence of PtsN_UnP_, the RCK domain in contrast to the CorC domain, remained less mobile. We suggest that the relatively rigid RCK domain may act as a platform to orient the CorC domain to engage with the transporter module and exert its inhibitory effect, indirectly. In this scenario the RCK domains act as allosteric scaffolds.

It is interesting that YcgO is a representative of a large class of K^+^-specific CPA1 antiporters present in numerous bacterial species including some pathogenic bacteria (Extended Data Figure 2). It would be worthwhile exploring the general applicability of the conclusions from this study to these bacterial systems and whether these studies could facilitate strategies for potential species-specific therapeutic interventions. In summary, this study highlights the interplay between a regulatory phosphorelay and the multidomain K^+^/H^+^ antiporter YcgO and delineates the roles of its various domains and their regulatory interactions.

## Methods

### Microbiological, genetic and molecular biology procedures

These procedures are described in the Supplementary information section. They include the lists of *E. coli* K-12 strains, plasmid constructs, oligonucleotide primers (Supplementary Tables 1, 2, and 3 respectively), media used and methods employed for strain construction and recombineering. Media used are also described.

#### Expression and purification of YcgO

An exponential phase (*A*_600_ of 0.6–0.8) culture of C41(DE3) cultured in LB medium, bearing the plasmid pHYD5011, expressing a C-terminally 3xFLAG hexahistidine tagged YcgO (YcgO_FH_) was generated. IPTG (at 1 mM) was added to induce expression of YcgO_FH_, followed by incubation at 20°C for 16 to 18 hours. Cells were harvested by centrifugation at 5000 *g* for 25 minutes at 4°C. The pellet was resuspended in lysis buffer (20 mM HEPES-K, 200 mM KCl, pH 7), supplemented with 1 mM PMSF and 0.4 mg/mL lysozyme. Cells were lysed using high-pressure homogenization at 4°C. All the purification steps were performed at 4°C. Unlysed cells were removed by centrifugation at ∼ 8000 *g* for 30 minutes, and the membranes were pelleted using ultracentrifugation at 1,10,000 *g* for 90 minutes. The membrane pellet was resuspended in solubilization buffer containing 20 mM HEPES-K (pH 7.0), 200 mM KCl,1 mM PMSF, and 10 mM LMNG. The suspension was gently mixed using end-over-end nutation for 3 hours at 4°C. Solubilized membranes were further clarified by ultracentrifugation at 1,11,000 g for 1 hour. The supernatant was kept for binding with pre-equilibrated Ni-NTA beads for 3 hours at 4°C. The beads were washed with wash buffer (30 mM imidazole, 20 mM HEPES-K, 200 mM KCl, 0.1 mM LMNG) and eluted in the elution buffer (300 mM imidazole, 20 mM HEPES-K, 200 mM KCl, 0.1 mM LMNG). Freshly eluted protein was concentrated using a 30 kDa cut-off concentrator and purified further by size exclusion chromatography (SEC) using Superose 6 increase 10/300 GL column equilibrated with SEC buffer (20 mM HEPES-K pH 7.0, 200 mM KCl, 2% v/v glycerol, 0.1 mM LMNG). Prior to injection, the concentrated protein sample was spun at ∼20,000 g for 20 minutes at 4°C to remove aggregates.

#### Expression and Purification of unphosphorylated PtsN

A mid-exponential phase culture (*A*_600_ 0.4 to 0.5) of the *E. coli* strain, GJ21015, bearing the plasmid pHYD6580, grown in LB medium containing 35 μg/ml chloramphenicol at 37°C, was obtained. To this culture, IPTG (at 0.5 mM) in the presence of 0.2% D-glucose, was added to induce expression of a C-terminally hexahistidine tagged PtsN and the culture was incubated for an additional 4-5 hours at 37°C. Glucose was included to prevent phosphorylation of PtsN by the glucose phosphotransferase system^13^. Cells were harvested by centrifugation and washed once with resuspension buffer (20 mM HEPES-Na, 200 mM NaCl, pH 7.0), prior to resuspension in lysis buffer supplemented with 1 mM PMSF and 0.4 mg/ml lysozyme. Cell lysis was performed using high-pressure homogenization at 4°C, and unlysed cells and debris were removed by centrifugation at 18,000 g for 30 minutes at 4°C. The resulting supernatant was supplemented with 10 mM imidazole and incubated with pre-equilibrated Ni-NTA resin (washed in buffer as above containing 10 mM imidazole) for 4 hours at 4°C. The resin was then washed with 40–50 column volumes of buffer containing 30 mM imidazole, and bound PtsN_UnP_ was eluted using buffer (20 mM HEPES-Na, 200 mM NaCl, pH 7.0) containing 300 mM imidazole. Protein expression was performed in GJ21015 because this strain in addition to lacking PtsN also lacks PtsP, consequently in this background overexpressed PtsN is present as PtsN_UnP_^14,31^. For the purification of C-terminally hexahistidine tagged PtsN with the H73E substitution, the plasmid pHYD5009 was transformed into the strain GJ18835 and purified using the same procedure as the above. The eluted protein was concentrated using a 10 kDa cut-off concentrator to a final volume of ∼0.5 mL and clarified by centrifugation at 20,000 g for 30 minutes at 4°C. The concentrated sample was subsequently loaded onto a Superdex 75 10/300 GL column for SEC using SEC buffer (20 mM HEPES Na, 200 mM NaCl, pH 7, 2% (v/v) glycerol).

#### Cryo-EM sample preparation and data collection

YcgO_FH_ was purified using SEC and concentrated to 5 mg/ml, and 3 µl of the final sample was applied to freshly glow-discharged QUANTIFOIL Au 1.2/1.3 holey carbon grids. Grids were blotted for 4.5 s under 100% humidity at 16°C and plunge-frozen in liquid ethane using a Vitrobot Mark IV system (ThermoFisher). For preparation of grids used in data collection of the PtsN_UnP_-bound YcgO_FH_ structure, YcgO_FH_ was incubated with PtsN_UnP_ at a 1:5 molar ratio for 1 h prior to vitrification.

Cryo-EM datasets were collected on a Titan Krios G4 (Thermo Scientific) microscope operated at 300 keV, equipped with a BioContinuum Gatan K3 direct electron detector and an Ametek-Gatan energy filter at the national cryoEM facility at IIT Bombay. Data were acquired at a magnification of 105,000×, corresponding to a pixel size of 0.85 Å in counted super-resolution mode. Movies were recorded using EPU software with a defocus range of –0.5 to –2.5 µm. Additional data collection parameters and statistics are summarized in (Extended Data Table 1).

#### CryoEM data processing and structure refinement

The YcgO_FH_ structure was resolved using a data of comprising 5835 dose-fractionated movies that were collected in EPU with 2x binning in super resolution mode and each movie had a total electron dose of 45 e⁻/Å².Data was processed in Cryosparc v4^35^. An initial particle number of 4,711,840 were picked up followed by 2D classification to filter the particle numbers to 264,367. Abinitio refinement yielded a 3D class with a subset of particles at 120,341 that was subjected to homogenous refinement and non-uniform refinement^36^ that yielded a map at resolution of 3.4 Å (at 0.143 GSFSC). For the YcgO_FH_-PtsN_UnP_ complex, a total of 8,510 dose-fractionated movies were collected in EPU using 2× binning, corresponding to a total electron dose of 45.5 e⁻/Å². All datasets were processed in CryoSparc v4^35^. Movie frames were aligned using patch motion correction with a maximum alignment resolution of 5 Å, and dose weighting was applied. The contrast transfer function (CTF) parameters for each micrograph were estimated using Patch CTF estimation, with a minimum and maximum CTF fit resolution of 25 Å and 4 Å, respectively. After CTF evaluation, 8,335 micrographs were retained for downstream processing. Initial particle picking was performed using a blob picker, yielding 529,127 particles. These particles were subjected to an initial round of 2D classification, and selected 2D class averages were used to generate templates for template-based particle picking, resulting in 3,242,861 particles. Following extensive 2D classification, 479,138 particles were selected for ab initio 3D reconstruction into two classes, followed by multiple rounds of heterogeneous and homogeneous refinement. Particles contributing to the best-resolved class were subjected to non-uniform refinement with C2 symmetry applied, resulting in a final 3D reconstruction which was used for fit the YcgO model derived from AlphaFold and PtsN crystal structure (PDB id 1A6J). Model was fit into the density using COOT and real space refinement was performed using PHENIX.refine in the PHENIX software suite^37,38^.

#### FSEC analyses

Fluorescence-detection size-exclusion chromatography (FSEC) was performed using a Shimadzu HPLC system equipped with a RF20A fluorescence detector and a temperature controlled autosampler^39^. Purified YcgO_FH_ was mixed with PtsN_UnP_ at a 1:10 ratio and incubated at 4°C for 1 h. Samples were injected into a Superose 6 Increase 10/300 GL column at a flow rate of 0.4 ml/min in a mobile phase comprising 20mM HEPES-K pH 7.0, 100 mM KCl, 100 mM NaCl, 0.1 mM LMNG. The shift in FSEC profile was observed by comparing YcgO_FH_-PtsN_UnP_ complex and YcgO_FH_. The intrinsic tyrosine fluorescence was measured λ_Ex_ at 275 nm and λ_Em_ at 305 nm. Sample preparation, incubation, and all chromatographic parameters were kept constant between trials to enable direct comparison of chromatograms.

#### Everted Vesicle preparation and assay

Primary cultures of GJ22224 bearing plasmids expressing under the control of the L-arabinose inducible promoter C-terminally 3xFLAG tagged YcgO, (YcgO_WT_), its D162N substitution derivative YcgO_D162N_ and the vector pBAD24, were grown in 10 ml KML medium supplemented with 100 µg/ml ampicillin for 12 h at 37°C. These cultures were sub-cultured in 800 ml of ampicillin supplemented KML medium and incubated at 37°C. Expression of the indicated YcgO proteins was induced with 0.05 % (w/v) L-arabinose at a culture *A_600_* of 0.6 and incubated at 20°C for 8 hours. Cells were harvested by centrifugation at 5000 g for half an hour at 4°C. The cell pellet was washed with a buffer comprising 20 mM Tris-HCl pH 7.5, 140 mM choline chloride and 10% glycerol (Buffer A) and resuspended in 60 ml of the same buffer containing 0.2 mg/ml of lysozyme. Resuspended samples were transferred into 300-400 ml sterile bottles and incubated for 30 minutes at 30°C with slow shaking at 60 rpm. The samples were homogenised at 400 bar for 3-5 minutes at 4°C in a GeoSavi PandaPlus high pressure homogenizer. The lysate was incubated for 30 minutes at 30°C with 10 mM MgSO_4_ and 10 µg/ml of DNase I with shaking at 60 rpm. Unbroken cells were removed by centrifugation at 13000g for 10 minutes at 4°C. Inside-out membrane vesicles were then isolated by ultracentrifugation at 110000 *g* for 1hour at 4°C and resuspended in 2 ml of Buffer A on end-to-end nutation for 4 hours. Vesicle aliquots of 50 ul were prepared and flash frozen in liquid nitrogen and stored at −80°C, till further use.

### Everted Vesicle Assay

Kinetic assay was performed in a Fluoromax spectrophotometer (Horiba) using a 2 ml fluorescence cuvette with a magnetic bead for fast equilibration. A 50 μl vesicle aliquot was resuspended in 1.9 ml of (Buffer A) (20 mM Tris-HCl, pH 7.5, 10% glycerol, 140 mM choline chloride). A final concentration of 40 μM ACMA dye was added to the reaction. Prescence of a stable baseline for ACMA fluorescence (excitation λ_ex_:409nm, λ_ex_:474nm) was monitored followed by addition of 100 μM ATP to energize the vesicles leading to quenching of ACMA fluorescence that reaches a baseline and monitored for about 50s. This was followed by addition of different monovalent cations (as chloride salts) to the cuvette (KCl, RbCl, NaCl, LiCl) at concentrations of either 5 or 10 mM to check for dequenching of ACMA fluorescence, an indicator of, cation induced H^+^ flux. After 50s of addition of monovalent cations to the mix, the reaction was quenched using NH_4_Cl at 10 mM to dissipate the pH gradient.

#### Immunoblotting assay with preparations of everted vesicles

A 50 μl aliquot of everted vesicles prepared in a buffer A was thawed and diluted in the same buffer with 1x SDS Laemmli buffer. The mixture was sonicated using Soniprep150 ultrasonifier (with a 15s on:off cycle) following which the sample was loaded onto a 12% SDS-PAGE gel. After electrophoresis, proteins were transferred onto a PVDF membrane at 100V for 100 minutes on ice. The membrane was stained with Ponceau (0.1% in 10% acetic acid solution in water) to estimate equal transfer of loaded samples followed by destaining the membrane using PBS-T (0.1% Tween 20 in phosphate buffered saline). Post destaining the membrane was blocked in 5% skimmed milk prepared in PBS-T for one hour at room temperature and probed with anti-FLAG antibody (MFLG-45A-Z) at 1:10000 overnight at 4°C. The blot was washed thrice with PBST followed by incubation with horseradish peroxidase conjugated secondary antibody (A28177) at 1:10000 for one hour at room temperature. The blots were developed using Immobilon HRP substrate (Millipore) and visualized on a Bio-Rad ChemiDoc Imaging System.

#### Isothermal titration calorimetry of YcgO vs PtsN

A MicroCal PEAQ-ITC instrument was used to measure the binding thermodynamics between YcgO_FH_ (9.5 μM) and PtsN (190 μM). The two PtsN proteins used were PtsN_UnP_ and PtsN_H73E_. YcgO and the PtsN solutions were prepared in the same buffer (20mM HEPES-K pH 7.0, 100mM KCl, 100mM NaCl, 0.1mM LMNG) to avoid buffer mismatch. All solutions were degassed immediately prior to titration. The sample cell (∼240 µl nominal volume) was loaded with YcgO_FH_, and the syringe (≈70 µL nominal volume) was loaded with PtsN proteins at a concentration approximately 20-fold higher than that of YcgO_FH_. A total of 19 injections of PtsN proteins (1.5 µL per injection) were made into the cell containing YcgO_FH_ during each titration, while the reference cell contained water. The time delay between injections was adjusted to allow the baseline to return to steady state, and the stirring speed was set to 700 rpm to ensure thorough mixing. All experiments were performed at 288 K (15 °C). Raw heat data were integrated peak by peak and normalized per mole of injectant to obtain a plot of the observed enthalpy change per mole of PtsN versus the PtsN:YcgO molar ratio. The enthalpy of dilution of PtsN, measured by titrating PtsN into buffer alone, was subtracted from the raw data before analysis. The first injection, affected by diffusion-related artifacts, was included in the raw data but excluded from curve fitting. Data processing and thermodynamic fitting were performed using the Malvern PEAQ ITC analyses software.

### Molecular Simulations

All simulation boxes were generated using the CHARMM-GUI membrane builder module^40^. Briefly, coordinates modelled in the cryoEM volume of YcgO-PtsN complex described earlier were taken as initial PDB for either PtsN free YcgO or PtsN-bound YcgO in simulation boxes. Lipid composition in the upper and lower leaflet were kept almost the same (i.e., ∼3:1 POPE:POPG), except cardiolipin concentration was kept at 10% in the lower (cytoplasmic) leaflet and 0% in the upper (periplasmic) leaflet of the membrane. A final concentration of 150 mM of KCl was utilized in these boxes. Positional restraints were removed from the atoms in a step-wise fashion over the course of two NVT and four NPT equilibration runs at 310K and with anisotropic pressure coupling. C-rescale and V-rescale were used as barostat and thermostat respectively for both equilibration and production simulations. The simulations were run on GROMACS 2024.2 and 2024.3 engines^41,42^. Replicates were run by generating 5 random seeds for each simulation box setup. The simulations were analyzed by aligning the Cα atoms, exporting the trajectories as PDB, and parsing and plotting them through in-house python scripts. Water coordinates were not exported for most of the runs, except when solvent accessibility analysis was done in a subset of runs. RMSF comparisons and RMS distance calculations were done using the first 100ns of each run.

## Data Availability

The EM maps and structure coordinates have been deposited in the PDB. Raw data for the manuscript is uploaded along with the manuscript.

Simulations performed as part of this study are included in Open Science Framework (OSF) and can be viewed via the following link: https://osf.io/b8vpe/overview?view_only=588fb076fd744326a5ad1aaa78842ce0.

## Author Contributions

AS was involved in sample preparation, biochemical experiments involving ITC, everted vesicles, chromatography, data collection and analyses. AA optimised early stages of sample preparation, functional assays and performed molecular simulations and some aspects of structural analyses. YP prepared the stains and plasmids employed in the study and performed all the growth assays in this study under the guidance of AAS. VS was involved in performing vesicle assays, immunoblotting and related data analyses. AP organised the project and performed the cryoEM data processing, structure refinements and planned the execution of biochemical experiments with inputs from AAS. Manuscript was written by AP and AAS with inputs from all the authors.

## Acknowledgements

Research in the manuscript was supported by the Revati and Satya Nadham Atluri Chair funds to AP from the Indian Institute of Science (IISc) and partly through the Wellcome Trust/DBT India Alliance Senior Fellowship (IA/S/22/1/506242) awarded to AP. Funding support by core funds from BRIC-Centre for DNA Fingerprinting and Diagnostics (CDFD) and a grant from the Department of Biotechnology, Government of India (BT/PR35617/BRB/10/1836/2019) to A.A.S is also acknowledged. AS and AA were former students of the Integrated PhD program of IISc. YP is a former student of the PhD program at the BRIC-CDFD, Hyderabad, affiliated to the Regional Centre for Biotechnology, Faridabad, India. VS is a student of the Integrated PhD program at IISc. The authors acknowledge the ANRF National CryoEM facility at IIT Bombay (IPA/2020/000413) and the technical support provided by Harshada Malvi during data collection. We also express our acknowledgement for the Advanced centre for cryo-electron microscopy facility at IISc for screening and preliminary data collection. Infrastructure support provided by the BRIC-CDFD is acknowledged. AP acknowledges DST-FIST support to the facilities at Molecular Biophysics Unit, IISc. MD simulations were run on virtual machines supported by the Google Cloud Research Credits program with the award GCP19980904.

## Extended Data Figures

**Extended Data Figure 1. Sequence alignment of YcgO homologues**. To avoid sampling-dependent bias and resulting misalignment of sequences, a protocol similar to the one described in an earlier study^4^ was used to generate initial alignment profile for the study. To do this, seed MSA was generated on all the annotated members of CPA1 (TC 2.A.36) and CPA2 (TC 2.A.37) families from transporter classification database, which were aligned using clustal omega with default parameters (i.e., gap opening penalty of 10.0, and gap extension penalties of 0.1 and 0.2 for pairwise and multiple sequence alignment respectively). Residues vital for K^+^ coordination and CPA1 motif are highlighted with blue line or asterisks. Red dot indicates salt bridge forming residues that stabilise CPA motif. This profile HMM was used to align a subset of sequences, depicted here, rendered using JalView, and secondary structure and domain limits were drawn manually. Every sequence is labelled with its generic name and its Uniprot accession ID.

**Extended Data Figure 2: Multiple sequence alignments of YcgO and PtsN homologues**. a, Clustal-based multiple sequence alignment of YcgO homologues from diverse species in proteobacteria that carry homologues of the *ycgO* gene. Secondary structure assignment is provided for the protein with dimerization helices in gray and transport helices in purple. RCK and CorC domain regions are indicated as pale blue and cyan respectively. Blue shading indicates greater than 80% sequence identity among residues. b, Clustal-based MSQ of PtsN homologues from proteobacterial species and colored in blue for residues having greater than 80% sequence identity. Orange boxes indicate residues interacting between PtsN and RCK domain of YcgO. Purple boxes indicate residue interactions between PtsN and CorC/HlyC domain of YcgO. Red asterisk indicates the H73 phosphorylation site.

**Extended Data Figure 3:** Purification of YcgO_FH_ (YcgO) and PtsN_UnP_ (PtsN). a, Panel represents the size exclusion chromatography (SEC) profile of purified YcgO in detergent containing buffer (20 mM Hepes, pH 7.0, 200 mM KCl, 2% glycerol, 0.1 mM LMNG using a Superose 6 10/300 GL column. The dashed vertical lines represent the fractions used for grid freezing. B, SEC profile of purified PtsN on a Superdex 75 column using a buffer comprising 20 mM HEPES pH 7.0, 200 mM KCl, 2% glycerol. C, SDS-PAGE gel (12%) displaying purified fraction of YcgO on the left of the size standard ladder (L) and PtsN at the bottom of the gel on the right.

**Extended Data Figure 4: Workflow of data processing and refinement steps for the PtsN free YcgO.** A dataset of 5385 movies was collected for this data which was curated down to 5345 micrographs. Scale bar for the representative micrograph depicts 50 nm. Initial particle picks yielded an original particle set of 4,711,840 particles that were subjected to repeated 2D classification resulting a set of 67389 particles that were reduced to 264367 particles that were subjected to *ab initio* refinement that yielded a 3D class with visible cytosolic domains with 120,321 particles. This set was subjected to homogenous refinement with C2 symmetry followed by non-uniform refinement to yield a map refined to 3.4 Å at a GSFSC cutoff of 0.143.

**Extended data Figure 5: Workflow of structure refinement for the YcgO-PtsN complex resolved through single particle cryoEM**. A total set of 8510 micrographs were collected as described in ED table 1. Scale bar in the representative micrograph represents 50 nm. After patch motion correction and CTF estimation the number of micrographs were curated down to 8335. Initial particle numbers of 3,814,921 were iteratively subjected to reference free 2D classification and Abinitio refinement yielding two classes which were combined to perform homogenous and non-uniform refinement yielding a map with a resolution of 3.2 Å.

**Extended Data Figure 6. Molecular Interfaces with the YcgO-PtsN_UnP_ complex**. The interfaces observed between the different domains of YcgO and its interactions with PtsN_UnP_ (PtsN) are indicated here. Interfaces include a, YcgO-PtsN complex structure with colored boxes indicating diverse interface observed between b, dimeric interface between YcgO TM helices mediated by phospholipid (yellow). c, RCK domains forming dimeric interactions d, PtsN interacting with RCK-CorC domain interface and e, CorC domain interacting with TM helices involved in elevator movement.

**Extended Data Figure 7. Structural overlaps of PtsN free YcgO with CPA transporters.** a, YcgO overlapped with KefC (PDB id 8BXG) yields a rmsd of 3.4 Å for 304 Cα atoms. b, Overlap with MjNhaP1 (PDB id. 4CZB) yields a rmsd of 2.13 Å for 278 Cα atoms. c, Overlap with EcNhaA (PDB id. 7S24) yields a rmsd of 11.0 Å for 251 Cα atoms. d, Overlap of YcgO with EcNapA which is in the outward-open state (PDB ID. 4BWZ) has a rmsd of 7.06 Å for 338 Cα atoms. As per the alignment values YcgO TM helices have the closest overlap within helix positions with MjNhaP. Overlaps were performed using molecular alignment option in Pymol. (www.pymol.org).e, Overlaps of KefC and YcgO protomer to display displacement of TM helices towards the cytosolic face. TMs 3, 10,13 display shifts closer to the vestibule in comparison to KefC. Disordered linker of TM12 that coordinates K^+^ ions is also displaced compared to KefC in YcgO.

**Extended Data Figure 8. Structural comparisons of RCK and CorC domains**. a, Overlap of an RCK domain (PDB id. 5F29) bound to cyclic-di-AMP (yellow) from SaCpaA transporter (salmon pink) with RCK domain of YcgO (slate) with a rmsd of 1.8 Å for 68 Ca atoms. b, Overlap of KefC RCK domain (PDB id. 8BXG) protomer (slate) over RCK domain of YcgO with a rmsd of 3.9 Å for 26 Cα atoms indicating high dissimilarity between the two domains. c, Overlap of CorC/HlyC domains from YcgO (deep blue) with similar domains from Nitrosomonas sp. (PDB id. 2P13; Cα rmsd = 1.57 Å), d, Neisseria meningitidis (2O3G; Cα rmsd = 1.37Å) and e, a domain from hemolycin C model from *Treponema hyodysenteriae* (Uniprot Q54318; Cα rmsd = 1.12 Å).

**Extended Data Figure 9**. **Simulation of domain flexibilities. a,** Cα trace of YcgO in complex with PtsN (left) (represents unphosphorylated PtsN) or in PtsN-free form (right). Root mean squared fluctuations over the course of a representative MD simulation trajectory is displayed as computed B-factor, going from low to high as variable widths and a colour gradient from blue to yellow. The B-factors representation were normalized to a common value for YcgO atoms in both the setups, and hence only a trace of PtsN is shown. **b,** Integrated-average of domain-wise RMSF in both the simulation setups. Data shown are for setups in which PtsN did not dissociate from the complex. Indicated *p*-values to assess domain flexibilities were calculated using Mann-Whitney U test.

**Extended Data Figure 10. Effect of water permeation in the vestibule of YcgO**. **a-c** MD simulations of YcgO in PtsN_UnP_ free form or in complex with PtsN_UnP_. Time-averaged projection of water molecules’ occupancy as heatmap with YcgO’s Ca trace and lipids overlaid for spatial reference. Representative frames from one out of five replicates for each setup are shown. Simulations were done with **a,** YcgO in complex with PtsN_UnP_ and **b,** YcgO in PtsN UnP free form, simulated in the presence of KCl. Only the first 100ns of simulation trajectories were processed to generate water map to avoid sampling conformational changes in YcgO. **c,** Inset from panel **c,** with spheres depicting residues S139, D272 and W276 in the water accessible cavity. **d**, Functionality of YcgO bearing the indicated substitutions. Cultures of the Δ*ptsN* Δ*ycgO* strain GJ22206 bearing the empty vector (pTrc99A) or pTrc99A expressing a functional N-terminal 3x FLAG tagged YcgO and its indicated substitution derivatives, were serially diluted and spotted on K_1_ and K_115_ glucose minimal agar plates containing 10 µM IPTG. Absence of growth in K_115_ medium is indicative of a functional YcgO. In the row marked vector growth is seen across dilutions in both K_1_ and K_115_ agar plates because the Δ*ycgO* mutation suppresses the K^+^ limited growth defect of the Δ*ptsN* mutation in K_115_ agar.

