## Extended Data figures and Supplementary figures for "Structural basis of K^+^/H^+^ antiport in YcgO and its inhibition by unphosphorylated PtsN"

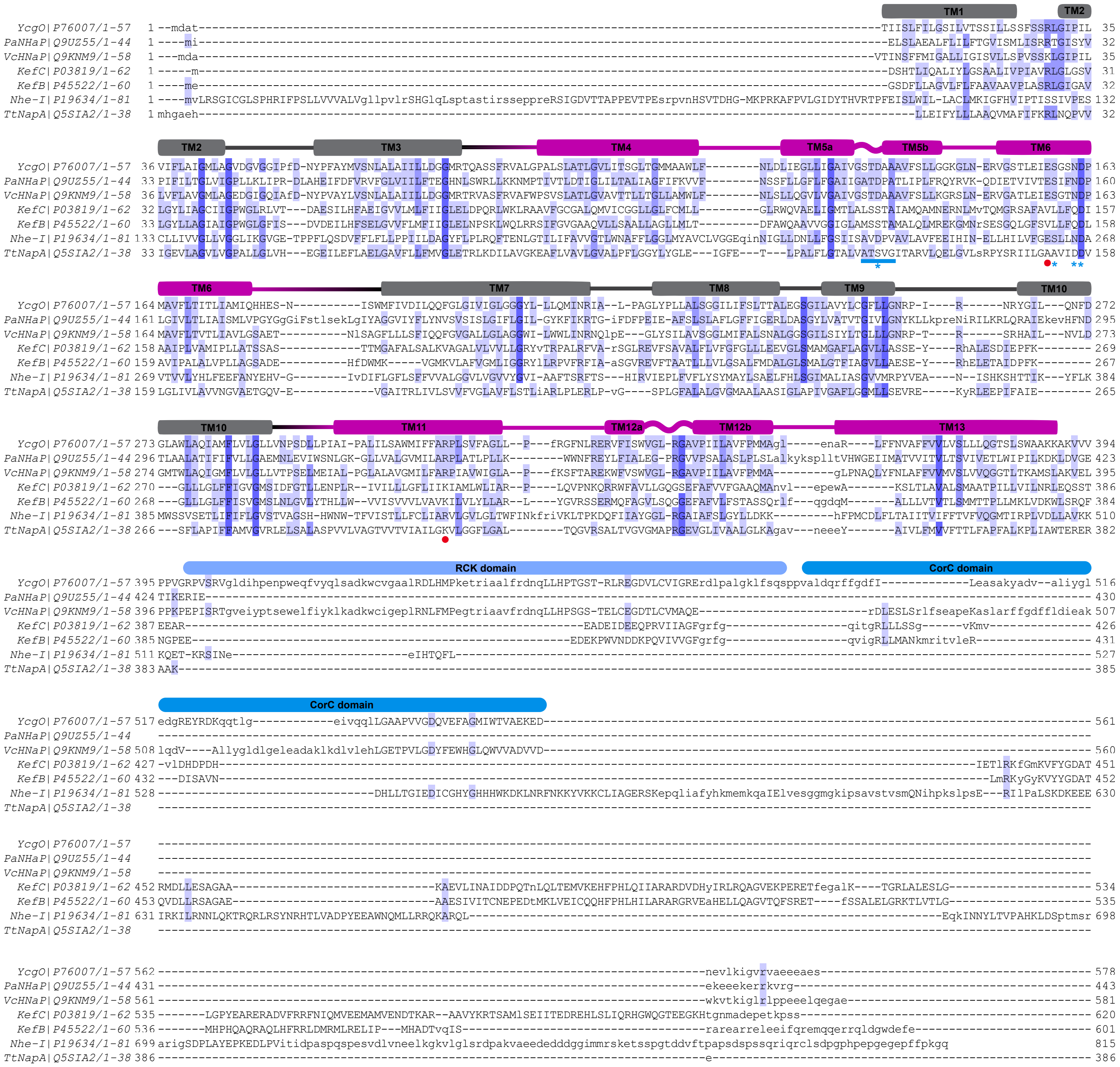


**Extended Data Figure 1. Sequence alignment of YcgO homologues.**

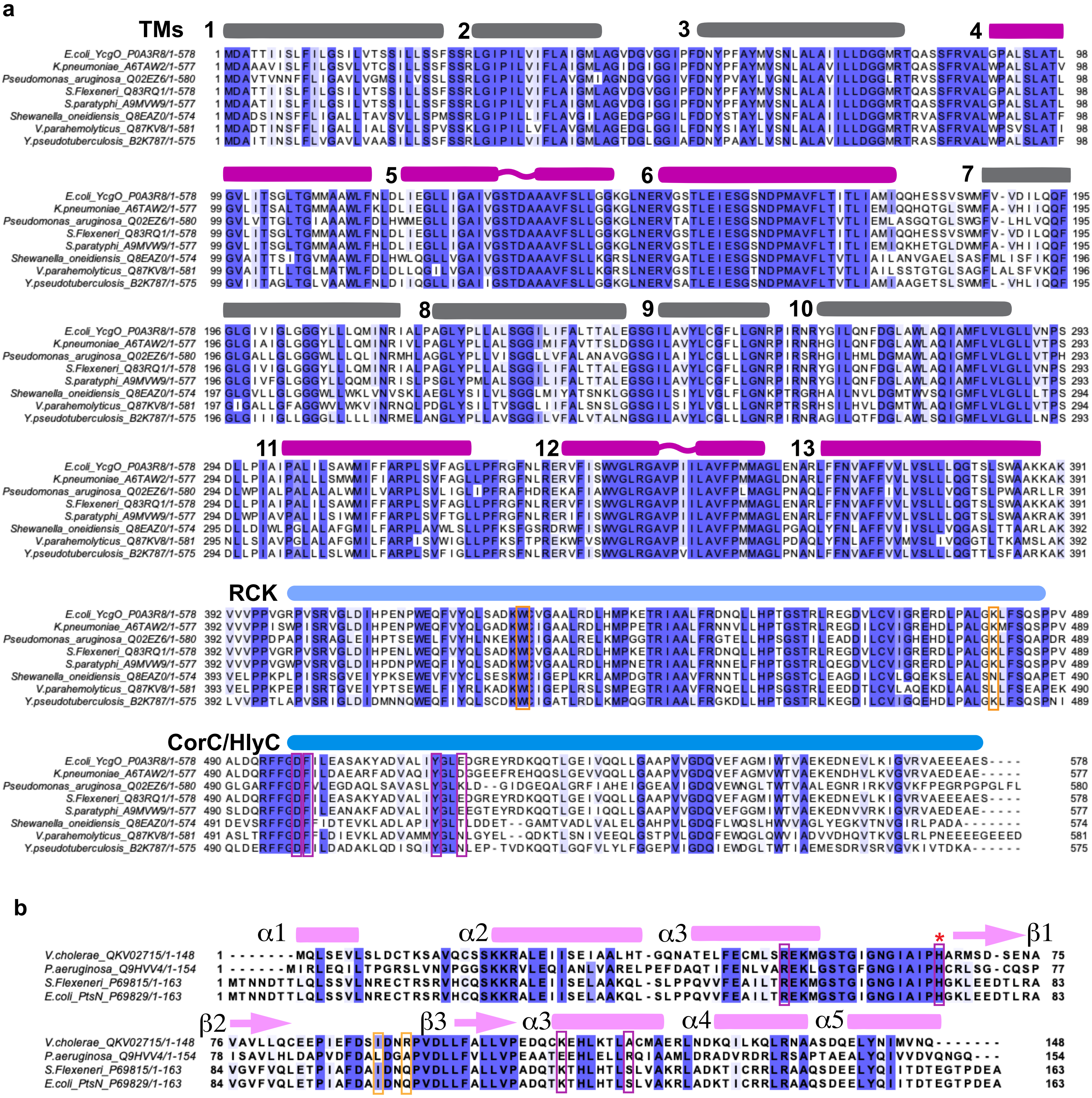


**Extended Data Figure 2: Multiple sequence alignments of YcgO and PtsN homologues.**

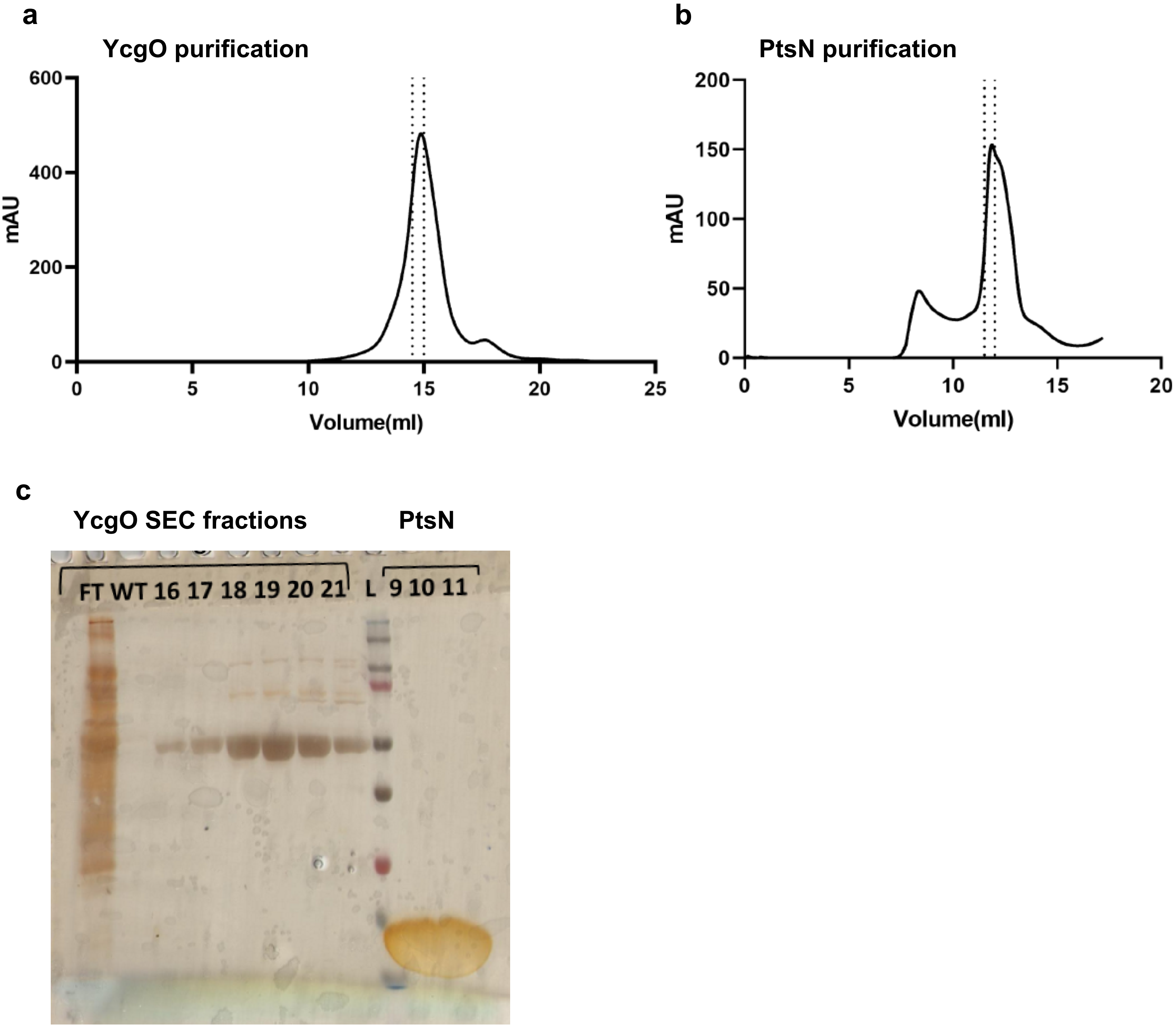


**ED figure 3: Purification of $\text{YcgO}_{FH}$ (YcgO) and $\text{PtsN}_{UnP}$ (PtsN) proteins.**

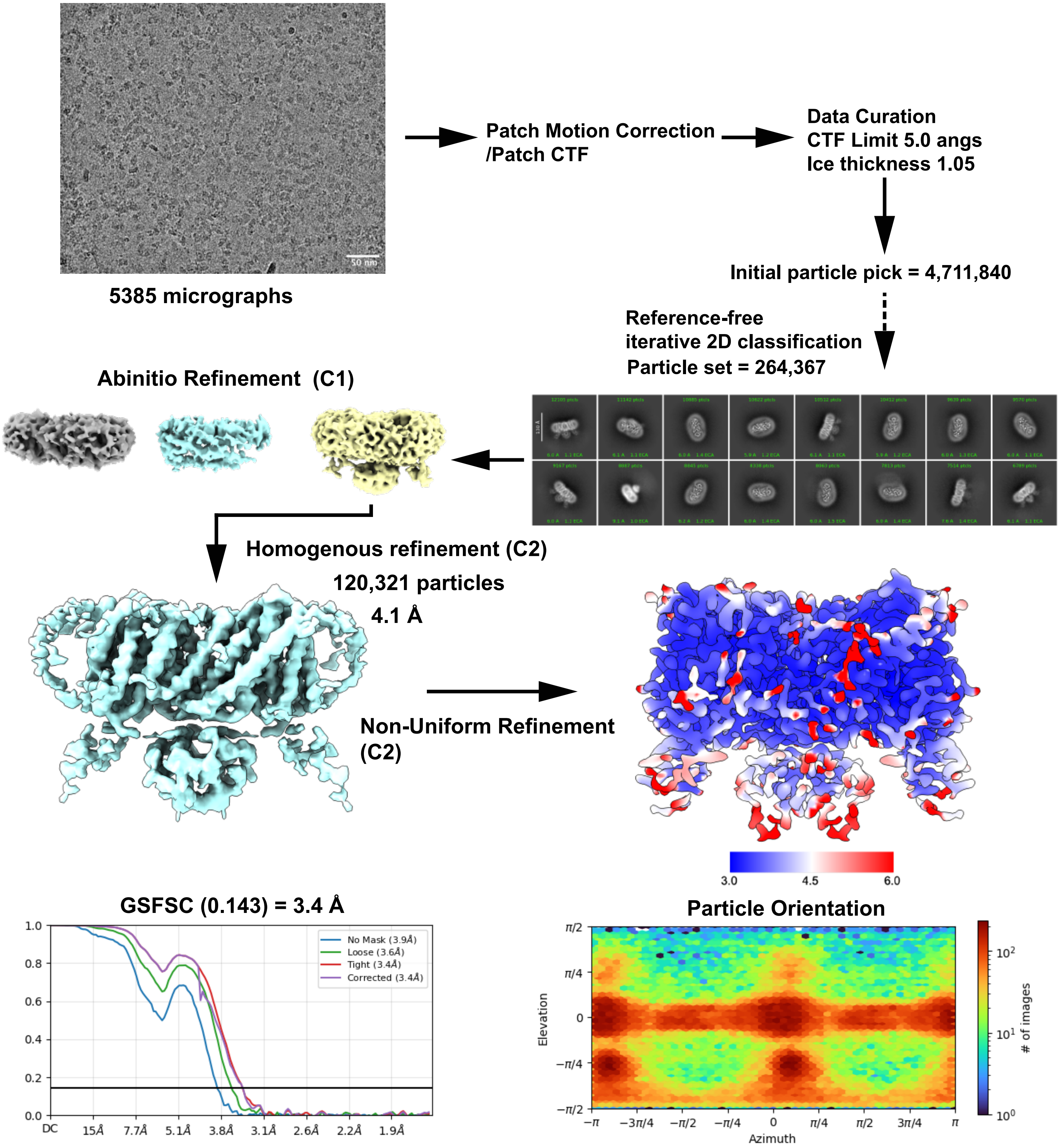


**Extended Data Figure 4: Work flow of data processing and refinement steps for the PtsN free YcgO.**

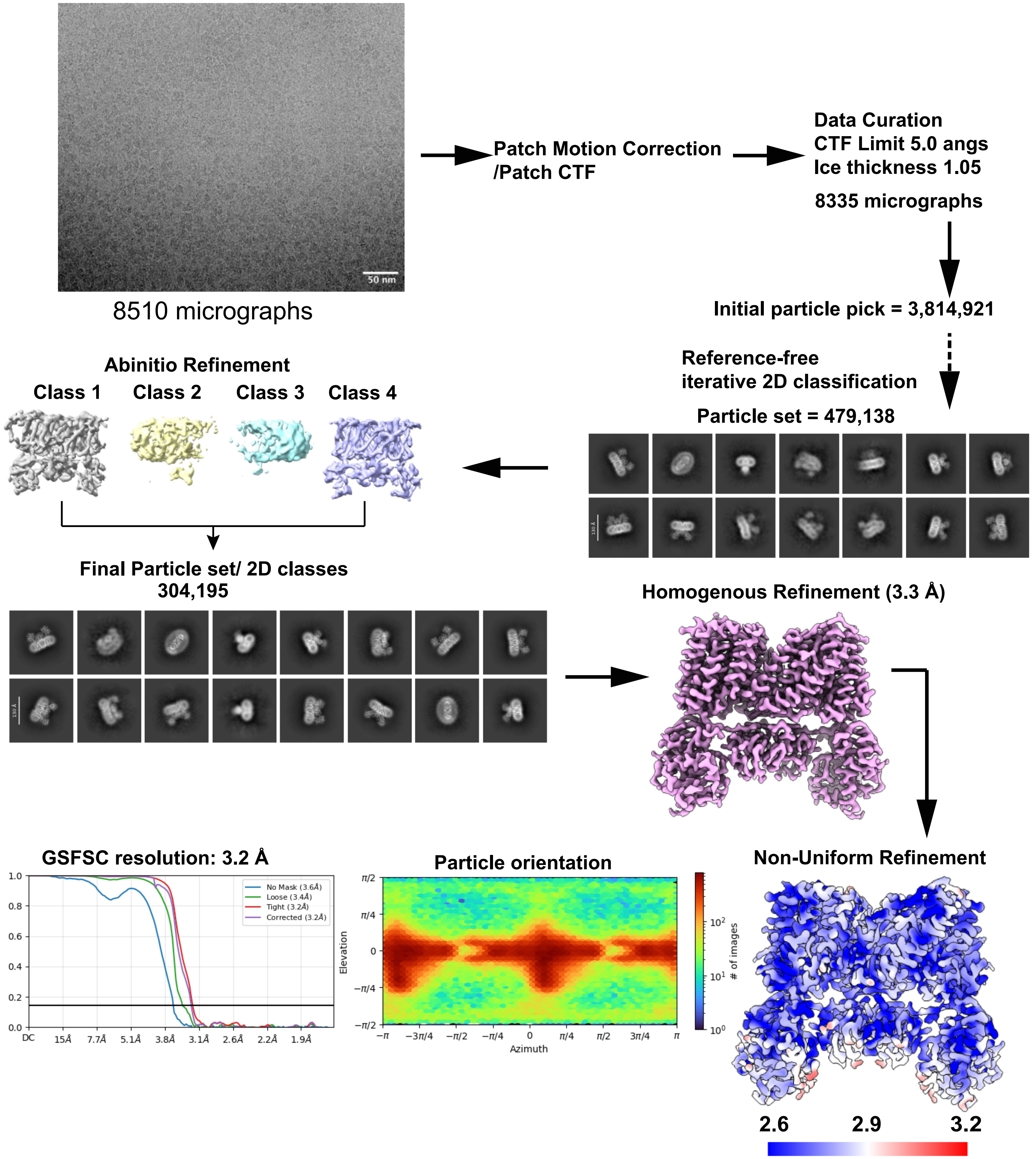


**Extended Data Figure 5. Workflow of structure refinement for the YcgO-PtsN complex resolved through single particle cryoEM.**

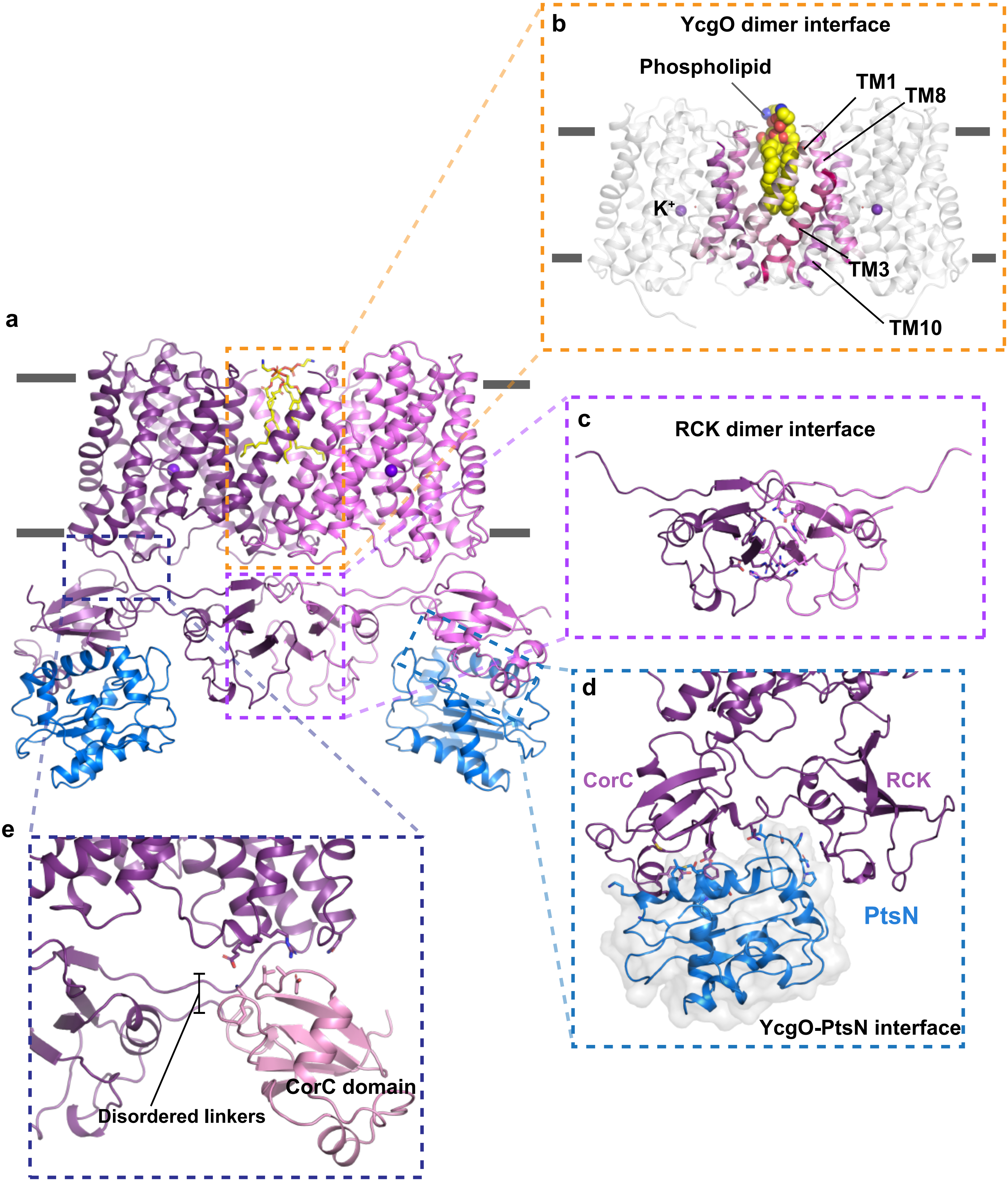


**Extended Data Figure 6: Molecular interfaces observed in the YcgO-PtsN$_{UnP}$ complex.**

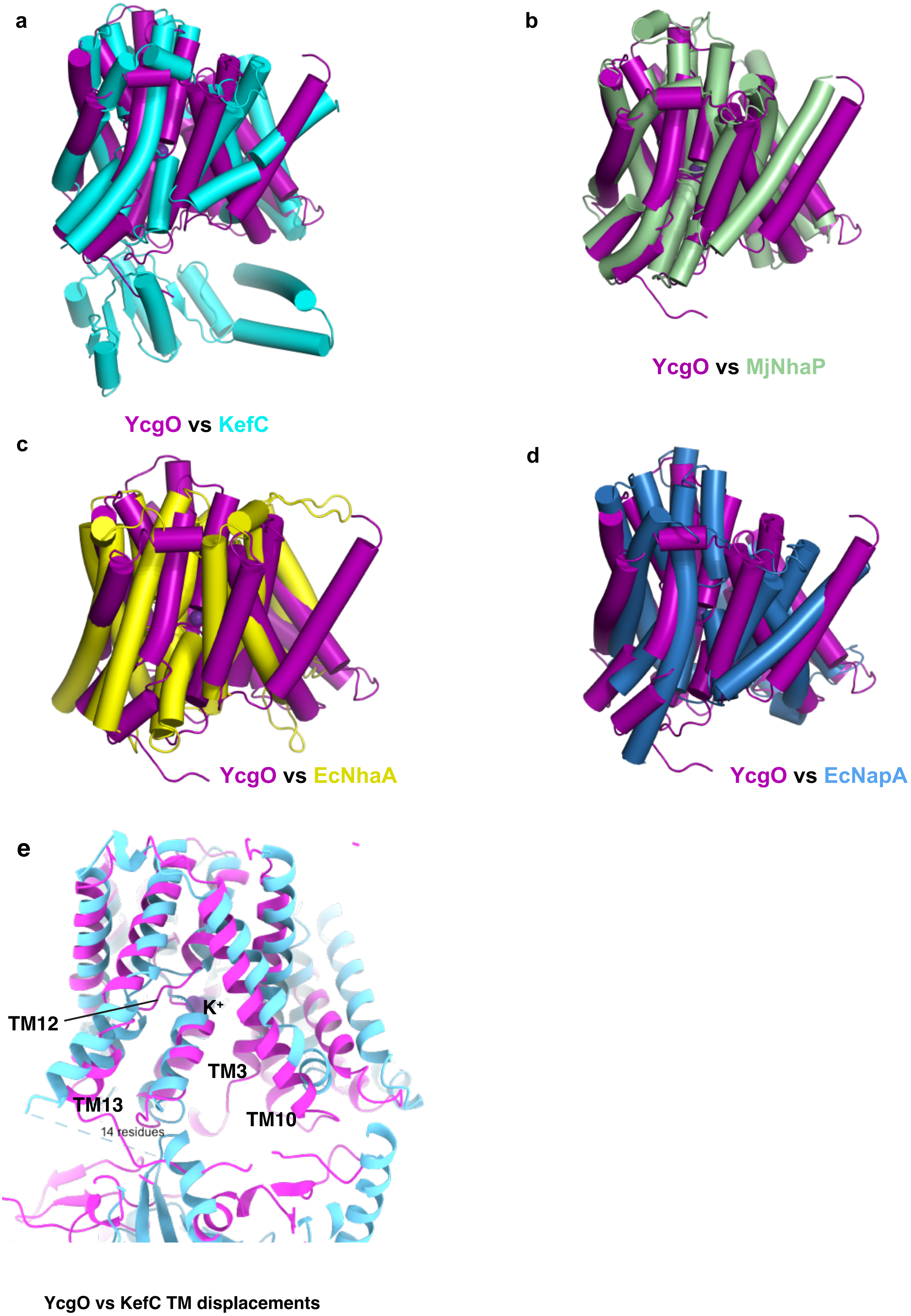


**Extended Data Figure 7: Structural overlaps of PtsN free YcgO with CPA transporters.**

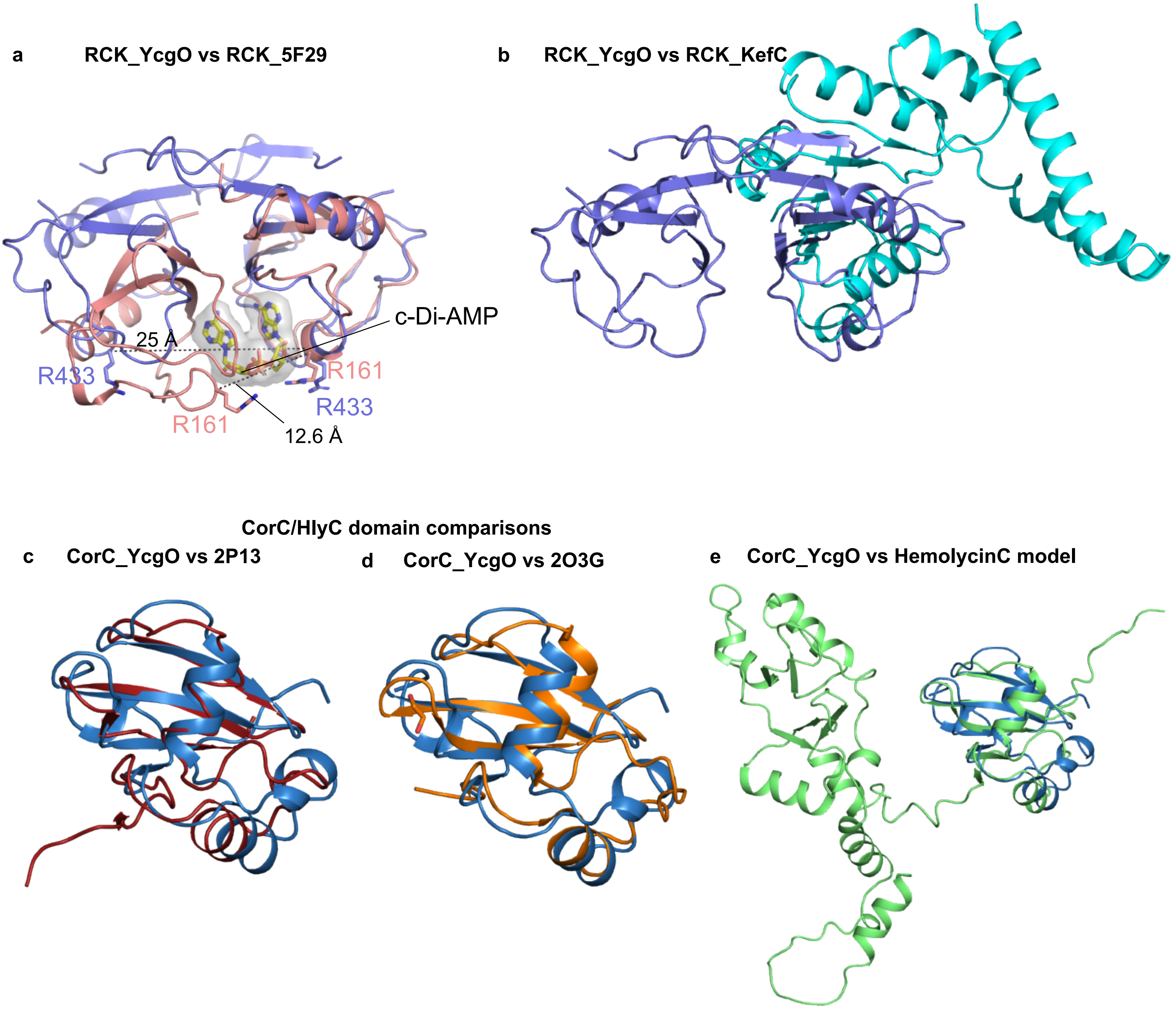


**Extended Data Figure 8. Structural comparisons of RCK and CorC domains.**

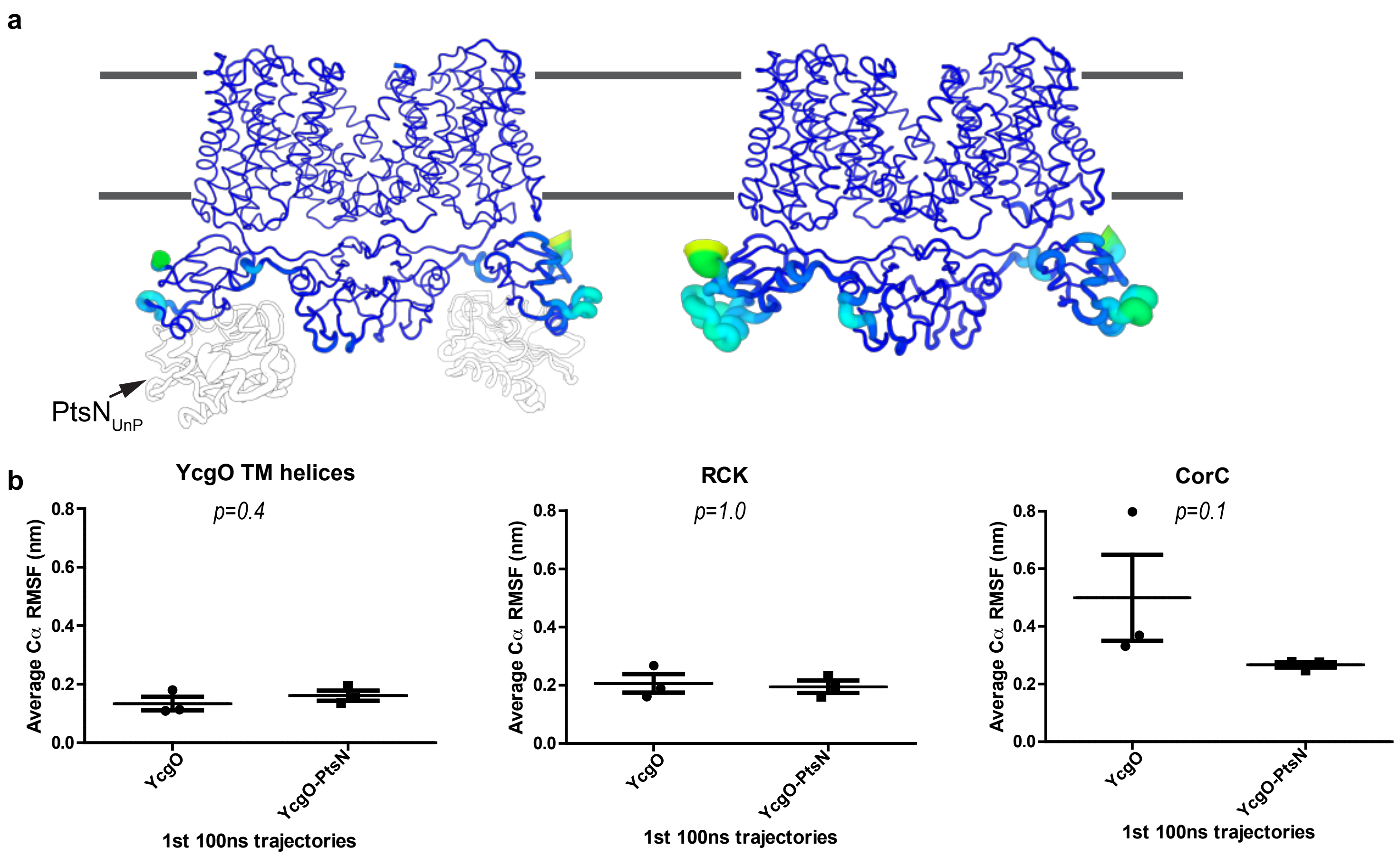


**Extended Data Figure 9: Simulation of domain flexibilities.**

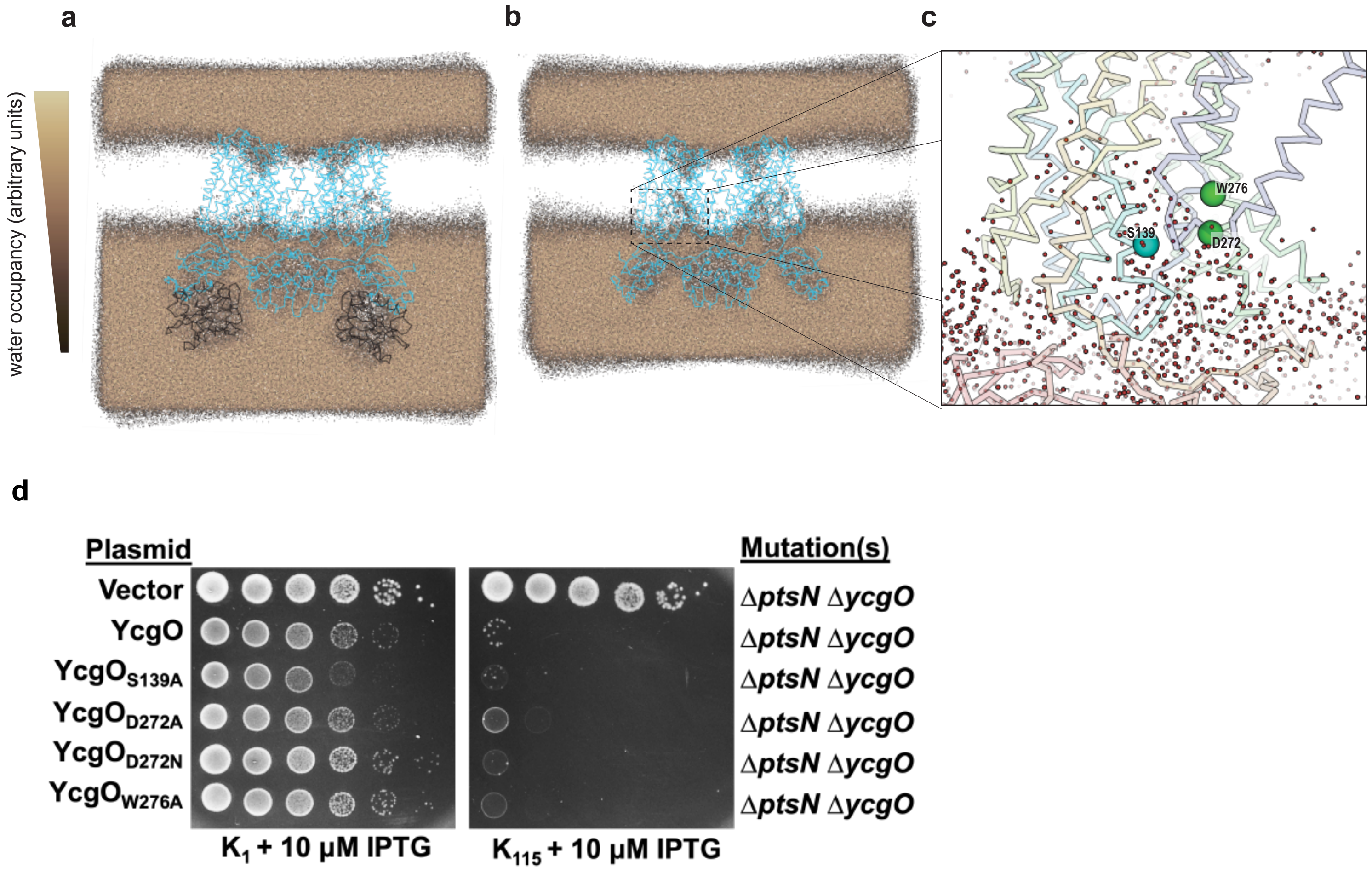


**Extended Data Figure 10. Effect of water permeation in the vestibule of YcgO.**

**Extended Data Table 1: Data Collection and Refinement statistics**

| Parameters | YcgO (PDB id.26OQ) | YcgO-PtsN (PDB id. 26PG) |
|---|---|---|
| **Data Collection** | | |
| **Microscope** | Titan Krios G4 | Titan Krios G4 |
| **Operating voltage** | 300 keV | 300 keV |
| **Magnification** | 105,000 x | 105, 000 x |
| **Detection** | Counted superresolution | Counted superresolution |
| **Energy filter (eV)** | 20 | 20 |
| **Defocus range** | -0.5 to 2.5 µ | -0.5 to 2.5 µ |
| **Angstrom (Å)/pixel (apix)** | 0.85 | 0.85 |
| **Total exposure (e/Å$^2$)** | 45.0 | 45.5 |
| **Spherical aberration (mm)** | 2.7 | 2.7 |
| **Refinement** | | |
| **Protein chains** | 2 | 4 |
| **Residues** | 795 | 1453 |
| **Ions** | 2 | 2 |
| **lipid** | 2 | 2 |
| **Total Number of atoms** | 6192 | 11366 |
| **B-factor (Å$^2$)** | | |
| **Protein** | 46.2 | 60.8 |
| **Lipid** | 70.7 | 66.5 |
| **water** | 39.8 | 48.8 |
| **RMS. Bond lengths** | 0.003 | 0.002 |
| **RMS Bond angles** | 0.57 | 0.41 |
| **RSCC map** | 0.87 | 0.84 |
| **GSFSC (0.143 cutoff)** | 3.4 | 3.2 |
| **Molprobity score** | 2.4 | 2.34 |
| **Clashscore** | 13.0 | 14.0 |
| **Ramachandran plot (%)** | | |
| **Favoured** | 94.1 | 94.7 |
| **Allowed** | 5.9 | 5.3 |
| **Outliers** | 0 | 0 |

### Structural basis of $K^+/H^+$ antiport in YcgO and its inhibition by unphosphorylated PtsN

Ayushi Srivastava[#,1], Arunabh Athreya[#,1,3], Yogesh Patidar[2,4], Vijeta Singh[1], Abhijit A. Sardesai[*,2] and Aravind Penmatsa[*,1]

**Supplementary Information: Table of Contents.**

I. Supplementary Figures.

1. S. Figure 1. Densities of TM helices of PtsN free YcgO.
2. S. Figure 2. Immunoblot of everted vesicles.
3. S. Figure 3. Alleviation of $K^+$ limited growth in K115 medium effected by $YcgO_{FH}$ in the absence of PtsN, by heterologous expression of Kup K+ uptake transporter.
4. S. Figure 4. Densities of TM helices and cytosolic domains in YcgO-PtsN complex

II. Supplementary Methods.
III. Supplementary Tables 1-3
IV. Supplementary References

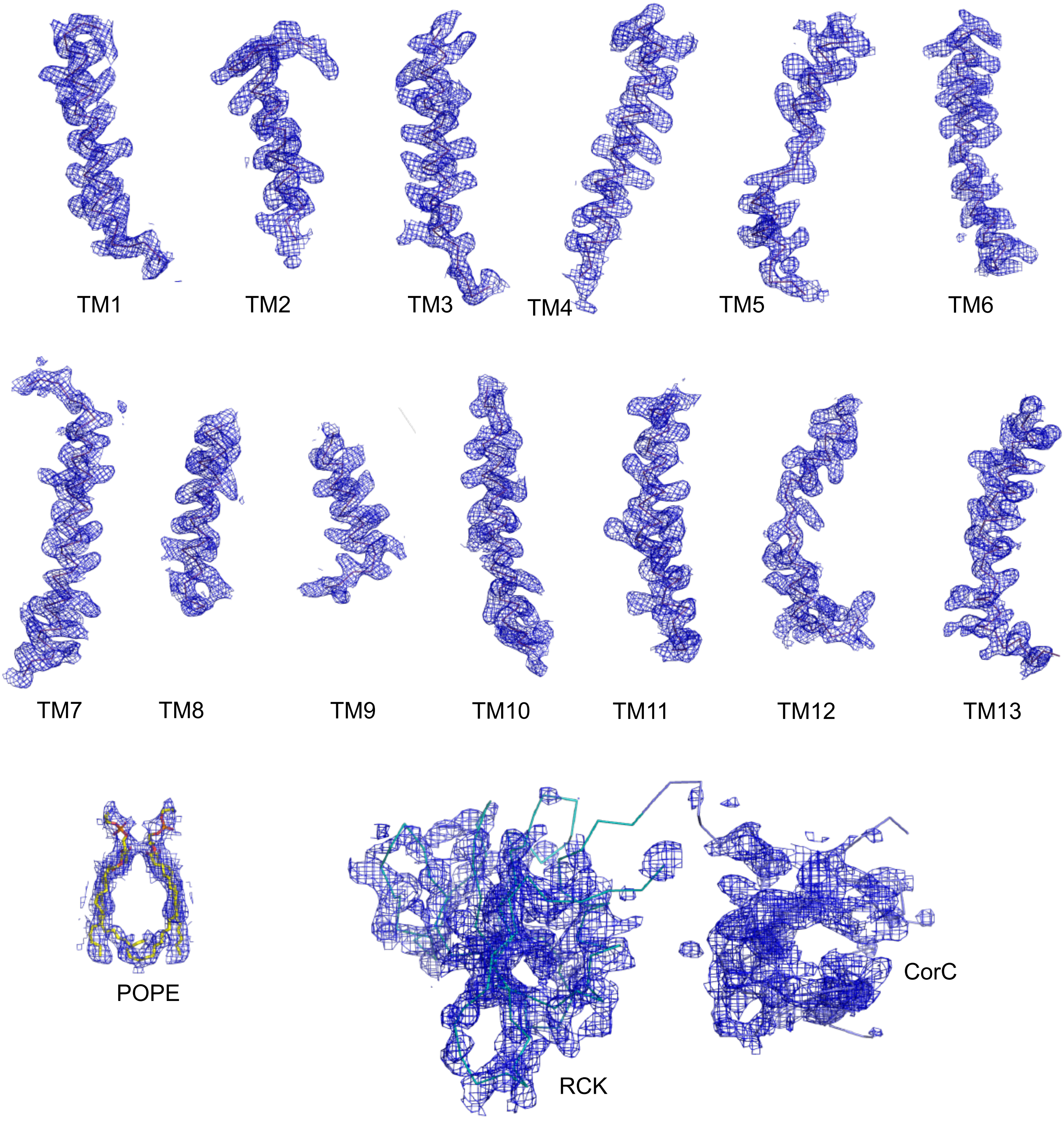


**Supplementary Figure 1.** CryoEM densities (blue) of individual transmembrane helices (purple) of YcgO and the cytosolic domains RCK (cyan) and CorC (slate). Densities (Blue) were contoured at a sigma level of 7.0 for TM helices (magenta), at 4.5 for POPE phospholipid (yellow). RCK and CorC domains were not modeled in the depositied structure but shown here with the weak densities observed in the map contoured at 6.0 sigma.

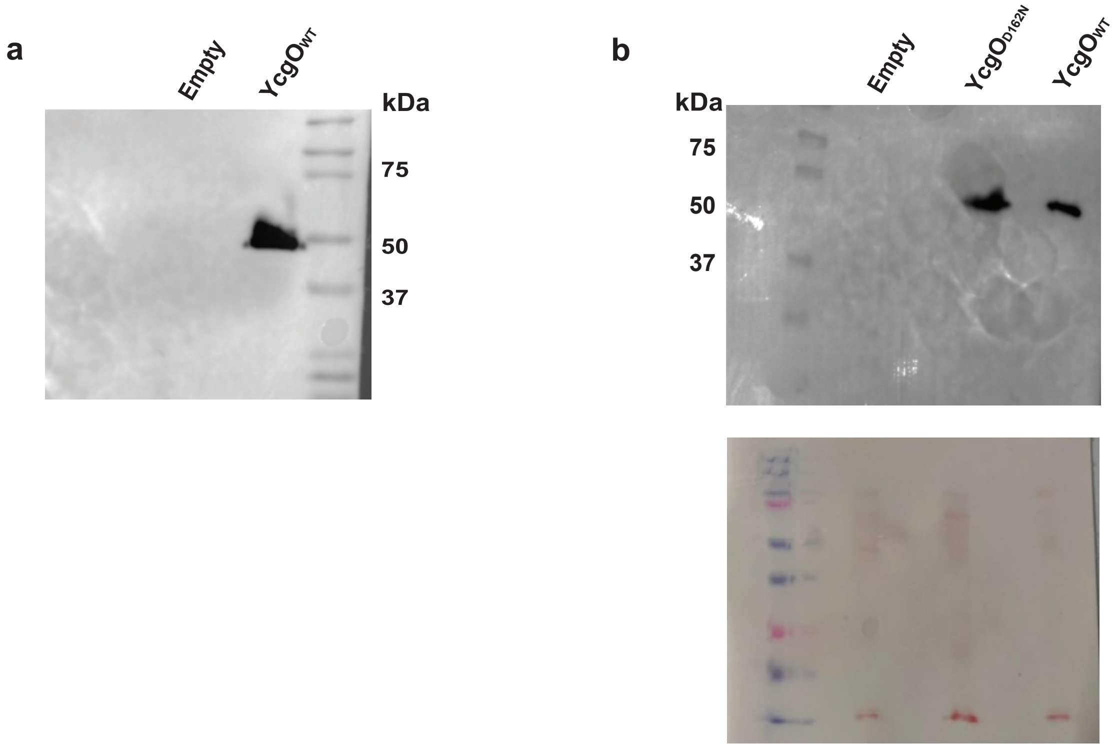


**Supplementary Figure 2:** a, Immunoblot of everted vesicles carrying $YcgO_{WT}$ and empty plasmids (PBAD24 in *E.coli* GJ22224) blottted with anti-FLAG antibody showing a band next to a Mol Wt of 50 kDa. b, Immunoblot of $YcgO_{WT}$ and $YcgO_{D162N}$ along with empty plasmid to demonstrate the expression of the protein in everted vesicles used to demonstrate $K^+$-induced $H^+$ flux. The blot was stained with Ponceau (0.1%) (below) to demonstrate equal loading of the vesicle fractions.

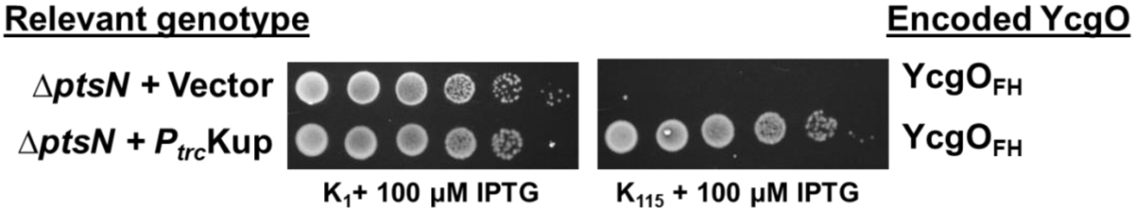


**Supplementary Figure 3**: **Alleviation of the $K^+$ limited growth in $K_{115}$ medium effected by $YcgO_{FH}$ in the absence of PtsN, by heterologous overexpression of the Kup $K^+$ uptake transporter.** Cultures of the strain GJ22302 that lacks PtsN (Δ*ptsN*) and expresses $YcgO_{FH}$ from its genomic location, bearing the vector and the plasmid pHYD1852 that expresses Kup from the IPTG inducible $P_{trc}$ promoter ($P_{trc}$Kup) were serially diluted and spotted on $K_1$ and $K_{115}$ agar plates bearing 100 µM IPTG. Strain and plasmid details are described in the supplementary Table 1 and Table 2. Figure represents one of two independent biological repeats.

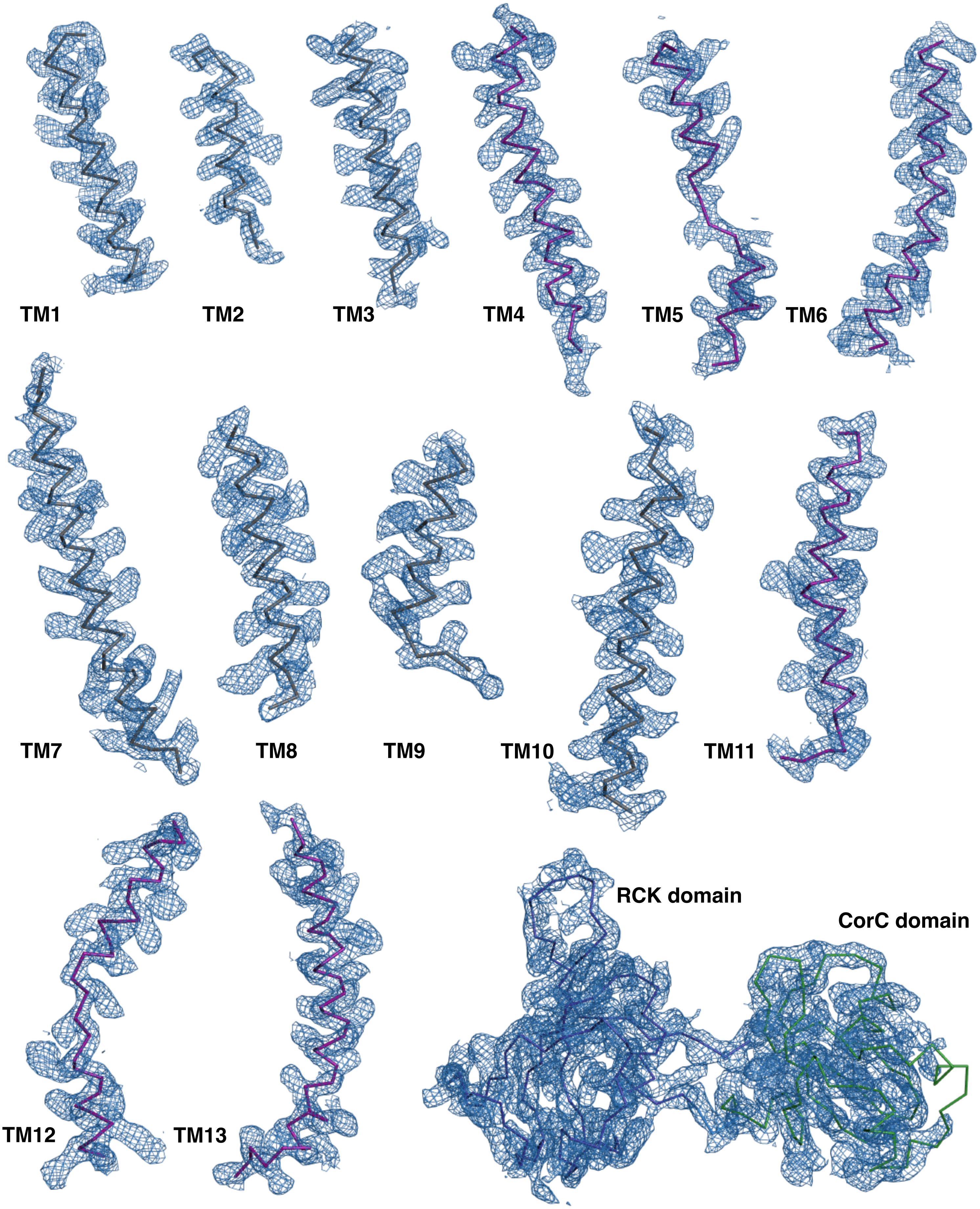


**Supplementary Figure 4:** CryoEM densities (blue) of individual transmembrane helices of YcgO and the cytosolic domains RCK (deep blue) and CorC (green). Densities were contoured at a sigma level of 6.0. Helices involved in dimerisation are in gray and the helices comprising transport/elevator domain are colored deep purple.

**Supplementary Methods.**

**Microbiological, genetic and molecular biology procedures**

Bacterial strains used in this study are derivatives of *E. coli* K-12 and their genotypes are listed in Supplementary Table 1. Strain construction was performed using P1*kc* (P1) transductions (Miller, 1992). Standard procedures for cloning, PCR and overlap extension PCR were employed. Deletion/insertion mutations of some genes were sourced from the Keio collection (Baba et al 2006) and details on these are described in supplementary Table 1. Plasmids and oligonucleotides used are listed in Supplementary Tables 2 and 3. Complex media used were LB agar/ broth and KML agar/ broth. KML is a derivative of LB medium in which NaCl is substituted by an equivalent concentration of KCl (Epstein and, 1971). Antibiotics were used at appropriate concentrations. IPTG (isopropyl β-D-thiogalactoside) and Ara (L-arabinose) inducers of the $P_{trc}$ and $P_{ara}$ promoters respectively, were used at the indicated concentrations and the growth temperature of bacterial strains was 37$^{O}$C. Two minimal media were used for assessing growth in the spot assays. They are 115 mM $K^+$ phosphate buffered medium and 115 mM $Na^+$ phosphate buffered medium, each of pH 7.2 (Epstein and Davies, 1970). $K_1$ medium represents 115 mM $Na^+$ phosphate medium containing 1 mM KCl and $K_{115}$ medium represents 115 mM $K^+$ phosphate buffered medium. These minimal media were supplemented with glucose (0.2%), $MgSO_4$ (1 mM) and agar as required.

**Recombineering procedures**

Recombineering (Datsenko and Wanner, 2000) was used to incorporate mutations or DNA encoding epitope tags in chromosomal *ycgO*. MG1655 expressing the wild-type C-terminally 3x FLAG-tagged YcgO (Sharma et al 2016) and its derivatives expressing the $YcgO_{D545A}$ and $YcgO_{\Delta CorC}$ versions were constructed by recombineering and are described in Patidar et al (2025, Table 1). For the construction of a chromosomally expressed C-terminally tagged derivative of YcgO, $YcgO_{FH}$ (YcgO appended with a 3x FLAG hexahistidine tag), the gene encoding 3x FLAG tagged *ycgO* was linked to the 6x His *kan* cassette of pSUB7 (Uzzau et al 2001) by overlap extension PCR, and the obtained PCR product was recombineered into MG1655 bearing the Δ*ycgO*::*cat* mutation (Patidar et al 2025) and the plasmid pKD46 as per Datsenko and Wanner (2000). Kanamycin (Kan) resistant ($Kan^R$) recombinants were isolated on Kan supplemented LB agar plates. All $Kan^R$ recombinants obtained were rendered chloramphenicol (Cm) sensitive, indicative of faithful recombineering events. A similar approach was used to construct a strain chromosomally expressing C-terminally 3x FLAG tagged derivative of YcgO, $YcgO_{R149A}$. In this case template DNA for PCR was a strain expressing a C-terminally 3x FLAG tagged derivative of YcgO with the required DNA alteration introduced into the targeting PCR amplicon via overlap extension PCR. The obtained genes were transferred to other strains by P1 transduction. Contribution of appended tags to YcgO is not considered for the amino acid numbering employed for proteins of this study, the natural numbering is used. Strains bearing genes encoding epitope-tagged proteins in the MG1655 background are not listed in Table 1. Primers used for recombineering are listed in Supplementary Table 3.

**Tests of growth phenotypes**

Dilution spotting was employed to assess growth phenotypes of strains used in this study. Overnight cultures of strains cultivated in an appropriate medium at 37˚C were normalized to an $A_{600}$ of 1 in $K_1$ medium, from where ten-fold serial dilutions were prepared in $K_1$ medium and 5 µl of each dilution was spotted on the surface of glucose supplemented $K_1$ and $K_{115}$ agar plates. IPTG and antibiotics when required were added as appropriate and plates were incubated at 37˚C for 16 to 24 hours. Representative images of dilution spotting performed in duplicates with independent cultures are shown.

**Immunoblotting Procedure**

For checking the expression of protein in everted vesicles, GJ22224 strain expressing pHYD6578(pBAD24-YcgO_3x FLAG) was employed. 50 μl of vesicles is mixed with 1x SDS Laemmli buffer and the reaction is diluted to a final volume of 500 μl in Buffer A (20 mM Tris-Cl (pH 7.5),140 mM Choline chloride and 10% glycerol) and sonicated using Soniprep150 ultrasonifier with 15 seconds on:off cycle. The protein sample is run in a 12% SDS-PAGE and transferred electrophoretically to PVDF membrane (Bio-Rad) using the wet transfer method at 100 V for 100 minutes on ice. The membrane is stained with 0.1% Ponceau S (Sigma) solution to confirm protein transfer. The membrane is then blocked using 5% skimmed milk prepared in PBS-T(Phosphate buffered saline + 0.1% Tween 20) for one hour at room temperature. After blocking the membrane is probed with anti-FLAG antibody (MFLG-45A-Z) (1:10000) overnight at 4°C and then washed thrice with PBS-T, followed by another incubation with horseradish peroxidase conjugated secondary antibody (Thermo Scientific A28177) (1:10000) for one hour at room temperature. The blot is washed with PBS-T and developed using chemiluminescent HRP substrate (Millipore) and visualized on a Bio-Rad ChemiDoc Imaging System.

**Supplementary Tables.**

**Supplementary Table 1: List of *E. coli* K-12 strains**

| Strain | Genotype | Comment/Feature |
|---|---|---|
| GJ17829 | MG1655 Δ*lacX74* Δ*trkA* Δ*kup* | Described in[a]. |
| GJ17831 | GJ17829 Δ*ptsN* | Described in[a]. |
| GJ17854 | GJ17829 *ycgO*:*kan* | Encodes a genomic C-terminal 3x FLAG-tagged YcgO, described in[a]. |
| GJ18835 | MG1655 Δ*lacX74* Δ*ptsN*::*kan* | Strain lacking PtsN, this study. |
| GJ19262 | GJ17829 Δ*ycgO*::*cat* | Described in[a]. |
| GJ19266 | GJ17829 $ycgO_{D545A}$:*kan* | Encodes C-terminal 3x FLAG-tagged YcgO bearing the D545A amino acid substitution, described in[a]. |
| GJ19291 | GJ17829 $ycgO_{\Delta CorC}$:*kan* | Encodes C-terminal 3x FLAG-tagged YcgO lacking CorC domain, described in[a]. |
| GJ21015 | MG1655 Δ*lacX74* Δ*ptsN* Δ*ptsP* | For purification of unphospho PtsN, described in[a]. |
| GJ22206 | GJ17831 Δ*ycgO*::*cat* | Described in[a]. |
| GJ22224 | MC4100 Δ(*araBAD*)567 Δ*ptsN* Δ*chaA* Δ*ybaL* Δ*ycgO* Δ*kefC* Δ*kefB* Δ*nhaA* | Described in[a]. |
| GJ22301 | GJ17829 $ycgO_{FH}$:*kan* | Encodes a genomic C-terminal 3x FLAG hexahistidine tagged YcgO. |
| GJ22302 | GJ17831 $ycgO_{FH}$:*kan* | Encodes a genomic C-terminal 3x FLAG hexahistidine tagged YcgO. |
| GJ22304 | GJ17829 $ycgO_{R149A}$:*kan* | Encodes a genomic C-terminal 3x FLAG-tagged YcgO bearing R149A amino acid substitution. |

[a]Patidar et al 2025; ":*kan*" indicates that the gene is linked at its 3' end to *kan* flanked by a pair of Flippase Recognition Target sites arranged in tandem.

**Supplementary Table 2: List of plasmids**

| Plasmid Name | Description |
|---|---|
| [a]pHYD5001 | [b]pTrc99A derivative, bearing a replacement of a unique NcoI site with an NdeI site. The pre-existing NdeI site in pTrc99A has been eliminated. |
| [a]pHYD5009 | pHYD5001 derivative, capable of expressing C-terminally hexahistidine tagged PtsN under the expression control of the IPTG inducible $P_{trc}$ promoter, bearing the H73E substitution. The corresponding gene is present in the NdeI and HindIII sites. |
| [a]pHYD1852 | Derivative of pHYD5001 with *kup* expression under $P_{trc}$ control and *kup* is present between EcoRI and HindIII sites. |
| [c]pHYD5011 | pTrc99A derivative, capable of expressing $YcgO_{FH}$, under $P_{trc}$ expression control. The corresponding gene is present in NcoI and HindIII sites and encodes YcgO, C-terminally abutted with a 3x FLAG hexahistidine tag. |
| [c]pHYD6578 | Derivative of [d]pBAD24, capable of expressing YcgO with a 3x FLAG tag appended to its C-terminus ($YcgO_{WT}$). The corresponding gene is present between the NcoI and HindIII sites. |
| pHYD6926 | pHYD6578 derivative, encoding YcgO in pHYD6578 bearing the D162N amino acid substitution ($YcgO_{D162N}$). |
| [c]pHYD6580 | Derivative of the plasmid pHYD5001 containing the *ori* and the *cat* gene from the plasmid [d]pBAD33, capable of expressing C-terminally hexahistidine tagged PtsN. The corresponding gene is placed between the NdeI and HindIII sites. |
| [c]pHYD6520 | pTrc99A derivative, encoding YcgO appended with an N-terminal 3x FLAG tag linked to its second amino acid. The corresponding gene is present between NcoI and HindIII sites. |
| pHYD6561 | pHYD6520 derivative, encoding YcgO in pHYD6520 bearing the D133N amino acid substitution. |
| pHYD6562 | pHYD6520 derivative, encoding YcgO in pHYD6520 bearing the E148Q amino acid substitution. |
| pHYD6563 | pHYD6520 derivative, encoding YcgO in pHYD6520 bearing the E155Q amino acid substitution. |

| | |
|---|---|
| pHYD6564 | pHYD6520 derivative, encoding YcgO in pHYD6520 bearing the E157Q amino acid substitution. |
| pHYD6565 | pHYD6520 derivative, encoding YcgO in pHYD6520 bearing the D162N amino acid substitution. |
| pHYD6912 | pHYD6520 derivative, encoding YcgO in pHYD6520 bearing the S139A amino acid substitution. |
| pHYD6913 | pHYD6520 derivative, encoding YcgO in pHYD6520 bearing the D272A amino acid substitution. |
| pHYD6914 | pHYD6520 derivative, encoding YcgO in pHYD6520 bearing the D272N amino acid substitution. |
| pHYD6915 | pHYD6520 derivative, encoding YcgO in pHYD6520 bearing the W276A amino acid substitution. |

Described in [a]Sharma et al 2016, [b]Amann et al 1988, [c]Patidar et al 2025 and [d]Guzman et al 1995.

**Supplementary Table 3: List of primers**

| Primer ID | Primer sequence (5'-3') | Purpose |
|---|---|---|
| JGAF_YcgO_EP_FP | TCTTAAAATGTAAAAAAATCACGCTTCAGGGCT | For the construction of chromosomal *ycgO*$_{FH}$ and *ycgO*$_{R149A}$ . |
| JGAF_YcgO_FH_FP | TCATGATATCGACTACAAAGATGACGACGATAAACACCACCATCATCACCAT | For the construction of *ycgO*$_{FH}$ on the chromosome. |
| JGAF_YcgO_FH_RP | ATGGTGATGATGGTGGTGTTTATCGTCGTCATCTTTGTAGTCGATATCATGA | As above |
| JGVYCGOF2 | TTCAGCAAAGCATCGCAACCGGCGATAGATGCTTCATTAAATTTACATATGAATATCCTCCTTAG | For the construction of *ycgO*$_{FH}$ and *ycgO*$_{R149A}$ on the chromosomal |
| JGAF_Y_R149A_FP | GGTAAGGGGCTTAACGAAGCGGTTGGCTCGACGCTGGAAA | For the construction of *ycgO*$_{R149A}$ on the chromosome |
| JGAF_Y_R149A_RP | TTTCCAGCGTCGAGCCAACCGCTTCGTTAAGCCCCTTACCA | As above |
| JGAF_Y_D133N_FP | CGCTATCGTCGGCTCTACCAATGCTGCAGCGGTCTTTTCT | For the construction of the plasmid pHYD6561 |
| JGAF_Y_D133N_RP | AGAAAAGACCGCTGCAGCATTGGTAGAGCCGACGATAG | As above |
| JGAF_Y_E148Q_FP | TGGTAAGGGGCTTAACCAGCGTGTTGGCTCGA | For the construction of the plasmid pHYD6562 |
| JGAF_Y_E148Q_RP | GCGTCGAGCCAACACGCTGGTTAAGCCCCTTAC | As above |
| JGAF_Y_E155Q_FP | TGTTGGCTCGACGCTGCAGATTGAATCCGGCAGTA | For the construction of the plasmid pHYD6563 |
| JGAF_Y_E155Q_RP | TACTGCCGGATTCAATCTGCAGCGTCGAGCCAACA | As above |
| JGAF_Y_E157Q_FP | CTCGACGCTGGAAATTCAGTCCGGCAGTAATGATC | For the construction of the plasmid pHYD6564 |

| | | |
|---|---|---|
| JGAF_Y_E157Q_RP | GATCATTACTGCCGGACTGAA TTTCCAGCGTCGAG | As above |
| JGAF_Y_D162N_FP | TTGAATCCGGCAGTAATAATC CAATGGCGGTCTTTCTGA | For the construction of the plasmid pHYD6565 |
| JGAF_Y_D162N_RP | AGACCGCCATTGGATTATTACT GCCGGATTCAATTTC | As above |
| JGAF_Y_S139A_FP | CGATGCTGCAGCGGTCTTTGC GCTGCTGGGTGGTAA | For the construction of the plasmid pHYD6912 |
| JGAF_Y_S139A_RP | CTTACCACCCAGCAGCGCAAA GACCGCTGCAGCATCG | As above |
| JGAF_Y_D272A_FP | GCATCCTGCAAAATTTCGCGG GCCTCGCCTGGCTGGCGCAA | For the construction of the plasmid pHYD6913 |
| JGAF_Y_D272A_RP | TTGCGCCAGCCAGGCGAGGC CCGCGAAATTTTGCAGGAT | As above |
| JGAF_Y_D272N_FP | CGGCATCCTGCAAAATTTCAA CGGCCTCGCCTGGCTGGCGC AA | For the construction of the plasmid pHYD6914 |
| JGAF_Y_D272N_RP | TTGCGCCAGCCAGGCGAGGC CGTTGAAATTTTGCAGGAT | As above |
| JGAF_Y_W276A_FP | CAAAATTTCGACGGCCTCGCC GCGCTGGCGCAAATCGCCATG TT | For the construction of the plasmid pHYD6915 |
| JGAF_Y_W276A_RP | AACATGGCGATTTGCGCCAGC GCGGCGAGGCCGTCGAAATTT TG | As above |
| D162N_FP | TCCGGCAGTAATAACCCAATG GCGGTCT | For the construction of the plasmid pHYD6926 |
| pBAD _RP | GCTTCTGCGTTCTGATTTAATC TG | As above |

Primers for sequence verification of the indicated plasmids and recombineering constructs are not listed. *ycgO*$_{FH}$ refers to the gene encoding the YcgO$_{FH}$ protein and *ycgO*$_{R149A}$ refers to the gene encoding a C-terminally 3x FLAG tagged YcgO with the R149A substitution.
